# RPA directly stimulates Mer3/HFM1 helicase processivity to ensure normal crossover formation in meiosis

**DOI:** 10.1101/2025.08.02.668043

**Authors:** Veronika Altmannova, Lucija Orlic, Carolina Carrasco, Céline Adam, Clara Aicart-Ramos, Dario Guerini, Valérie Borde, Joao Matos, Fernando Moreno-Herrero, John R. Weir

**Affiliations:** Friedrich Miescher Laboratory of the Max-Planck-Society, Max-Planck-Ring 9, 72076 Tübingen, Germany; Max Perutz Labs Vienna, Vienna BioCenter, Dr.-Bohr-Gasse 9, 1030 Vienna, Austria; Vienna BioCenter PhD Program, a Doctoral School of the University of Vienna and the Medical University of Vienna, A-1030 Vienna; Centro Nacional de Biotecnologia, CSIC, C/ Darwin 3, Campus UAM 28049, Cantoblanco, Madrid, Spain; Institut Curie, PSL University, Sorbonne Université, CNRS UMR3244, Dynamics of Genetic Information, 75005, Paris, France

**Keywords:** meiosis, RPA, DNA repair, helicases, crossovers

## Abstract

Meiotic crossover formation is critical for generating viable gametes and enhancing genetic diversity. The helicase Mer3 (HFM1 in humans) is a highly conserved factor essential for promoting crossovers and ensuring their proper distribution. Here, we identify replication protein A (RPA) as a direct interactor of budding yeast Mer3. We demonstrate that this interaction is conserved between human HFM1 and RPA. Cross-linking mass spectrometry and structural modelling with AlphaFold2 reveal a conserved and specific Mer3-RPA interface. Single-molecule magnetic tweezers assays demonstrate that direct RPA interaction is required for Mer3 helicase processivity under conditions of low DNA tension. Consistently, a *mer3* mutant deficient in RPA binding exhibit reduced crossover frequencies and accumulate unresolved recombination intermediates during budding yeast meiosis. Via genome-wide localisation experiments, we link this effect to a weakened recruitment to double-strand break sites of the mer3 mutant. Our findings provide mechanistic insights into coordination of meiotic recombination by the Mer3 helicase through interactions with the canonical DNA repair machinery, highlighting a conserved mechanism underlying crossover control during sexual reproduction.

## Introduction

The generation of viable, haploid gametes - eggs and sperm in mammals; spores in yeast - is essential for the continuation and evolution of eukaryotic life. Haploidization during gametogenesis depends on the segregation of homologous chromosomes during meiosis I. Faithful segregation of homologous chromosomes during the first meiotic division is indispensable for fertility and species survival. Crossovers (COs) between homologs create the physical connections that enable their bi-orientation on the meiotic spindle [1], yet the very same DNA exchanges are suppressed in somatic cells because they can provoke deleterious chromosomal rearrangements and loss-of-heterozygosity [2]. Understanding how meiotic cells generate COs efficiently while minimising collateral damage is therefore a central problem in chromosome biology, with ramifications for sexual reproduction, infertility and evolution.

Meiotic recombination is initiated by programmed DNA double-strand breaks (DSBs) created by the topoisomerase Spo11 [3,4]. Resection of meiotic DSB ends by MRX/Sae2 followed by the Exo1 pathway produces 3 single-stranded DNA (ssDNA) tails [5]. These overhangs are immediately coated by the heterotrimeric replication protein A (RPA), which in budding yeast consists of the large Rfa1 (*≈*70kDa), medium Rfa2 (*≈*32kDa) and small Rfa3 (*≈*14kDa) subunits [6,7]. Each subunit contributes one or more oligonucleotide/oligosaccharide-binding (OB) or winged-helix domains, which are connected by flexible linkers to form a dynamic architecture that can wrap 30nt of ssDNA [8]. Two surfaces within the RPA assembly are known to serve as versatile docking sites for partner proteins: the basic N-terminal OB-fold of Rfa1 (N-OB) [9,10] and the acidic, phosphorylation-rich C-tail of Rfa2 [7].

By masking ssDNA, RPA prevents secondary-structure formation, recruits the Mec1/ATR checkpoint kinase, and orchestrates downstream repair steps [11]. Mediator complexes and Rad51-paralogs modulate the association of RPA with ssDNA to allow Rad51 and Dmc1 to assemble onto ssDNA creating the presynaptic filament, enabling strand invasion and displacement-loop (D-loop) formation [12].

Several superfamily-2 (SF2)-type helicases sculpt these early intermediates. Sgs1/BLM, and Mph1/FANCM dismantle D-loops to bias repair toward non-crossover products, thereby safeguarding genome integrity [13–16]. Paradoxically these factors are also required for normal crossover formation during meiotic recombination, presumably due to their ability to recycle aberrant CO intermediates [17,18]. Proteins of the ZMM ensemble, Zip2-Zip4-Spo16, Msh4-Msh5 and the helicase Mer3, bias repair toward COs and enforce their spacing along chromosomes [19,20]. RPA is not a passive ssDNA coat: it can directly stimulate helicase motors. Human BLM and its yeast ortholog Sgs1 bind the Rfa1 N-OB motif, increasing unwinding processivity on kilobase duplexes [21,22]. Werner syndrome helicase (WRN) contacts the acidic RFA2 C-terminus, accelerating the resolution of G-quadruplex and forked structures [23,24].

Branch-migration motors FANCM and the annealing helicase SMARCAL1 are targeted to RPA-coated forks via phospho-dependent docking on RFA2, which enhances fork-regression efficiency [25]. In meiosis, how helicase activities are selectively tuned to promote or suppress CO formation remains poorly understood.

Mer3/HFM1 is a SF2 helicase, and is structurally related to RNA helicases including Brr2 [26] and Mtr4 [27], which contain a helicase domain, made up of two RecA-like domains, followed by an extended C-terminal domain (Figure 1A). Mer3 has been previously shown *in vitro* to stabilise D-loop repair intermediates and extend the D-loop, which may promote second end capture [28]. While the addition of RPA resulted in longer tracts of strand separation *in vitro*, this effect was considered to be due to the ssDNA binding properties of RPA, since bacterial SSB had the same effect on Mer3 [29].

**Figure 1.**
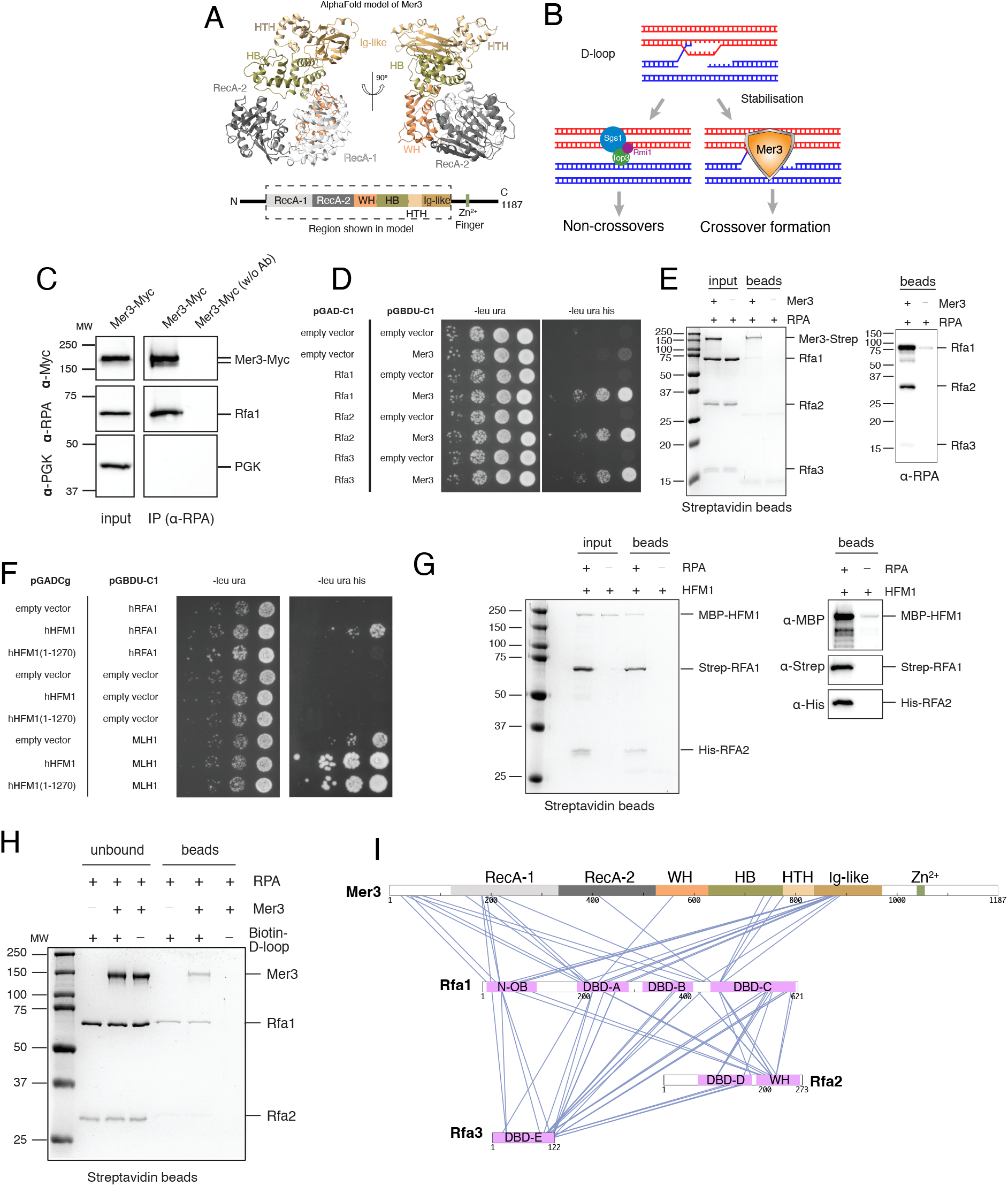
Mer3 helicase binds directly to RPA. A) Structural prediction (AlphaFold2) and domain cartoon of *S. cerevisiae* Mer3. For the AlphaFold prediction, only the structured region (dashed region of domain cartoon) is shown. B) Current model for the activity of Mer3 and its role in meiotic crossover formation. C) Co-IP *S. cerevisiae* cultures (SK1 strain, yWL514) after 5 hours in sporulation media. The same input samples (left panel) were used in the presence and absence of anti-RPA antibodies, revealing an association between Mer3 and RPA. D) Yeast two-hybrid assay of Mer3 with the three RPA subunits - Rfa1, Rfa2 and Rfa3. Empty vectors (either pGAD or pGBDU) are used as controls for non-specific activation. Specific association is shown between Mer3 and all three RPA subunits. E) Pulldown on streptavidin beads of recombinant C-terminally 2xStrepII-tagged Mer3 and RPA complex. Since only Rfa1 is clearly visible in the coomassie stained gel, we used anti-RPA antibodies to show the specific *in vitro* binding of all three subunits of RPA to Mer3 (inset, right). F) Yeast two-hybrid assay of human HFM1 (hHFM1) and human RFA1 (hRFA1). Full-length hHFM1 shows a specific association with hRFA1, whereas C-terminally truncated hHFM1 (amino acids 1-1270), does not. Nonetheless hHFM1 1-1270 does still interact with human MLH1. G) Coomassie stained SDS-PAGE of a pulldown on streptavidin beads of 2xStrep-II-tagged human RPA complex (RFA1 and RFA2 indicated) against MBP-HFM1. Western blotting (right) confirms the identity of the bands indicated, and the specificity of the pulldown, confirming that the interaction of HFM1 and RPA is conserved between budding yeast and humans. H) Coomassie stained SDS-PAGE of a pulldown of biotinylated D-loop DNA on streptavidin beads. Both RPA and the RPA-Mer3 complex seem to bind to D-loop DNA equally well. I) Cartoon representation of the inter-subunit cross-links from a cross-linking mass-spec (XL-MS) experiment on a Mer3-RPA complex using the DSBU cross-linker. Only cross-links between Mer3 and Rfa1 are observed.

Given RPA’s central placement at ssDNA at repair intermediates and that Mer3 helicase dead mutants resulted in phenotypes more mild than *mer3*Δ alleles [30,31], we reasoned that Mer3 might engage RPA directly to fine-tune its activity on recombination intermediates. We therefore set out to test whether Mer3/HFM1 physically interacts with RPA, to map the interface, and to define how this interaction influences helicase behaviour and meiotic crossover formation. Here we demonstrate that Mer3 binds the conserved Rfa1 N-OB cleft via a short C-terminal helix, that this contact enhances Mer3 processivity on DNA under low tension, and that weakening the interface perturbs crossover formation *in vivo*.

## Results

### Mer3 binds directly to the RPA complex in a conserved interaction

As described above, several genome-stability DNA helicases, including BLM/Sgs1 bind directly to RPA [21,22]. Mer3 works in part to antagonise Sgs1 through binding to some of the same binding partners including Top3 and Rmi1 [32]. Therefore we asked whether Mer3 might also bind directly to RPA. We found that Mer3 with a C-terminal 9xMyc tag (Mer3-Myc) interacts with Rfa1 in an anti-RPA co-immunoprecipitation experiment from meiotic *S. cerevisiae* (SK1 strain) cultures after 5 hours in sporulation media (Figure 1C), which is when meiotic repair intermediates are present. Furthermore, using an yeast two-hybrid assay (Y2H), we detected an interaction between Mer3 and all three RPA subunits (Rfa1, Rfa2, Rfa3) (Figure 1D), indicating that the Mer3-RPA interaction was independent of other meiosis-specific factors. We note that all three RPA subunits are well expressed in vegetative yeast (several thousand molecules of each subunit per cell [33]), therefore making it a realistic possibility that Mer3 binds directly to only one of the three subunits.

We considered that, since both RPA and Mer3 [32] can bind to ssDNA [34,35], a Mer3-RPA interaction could be mediated indirectly by ssDNA. To address this, we purified recombinant budding yeast RPA, expressed in bacteria, to homogeneity (Supplementary Figure 1E). We likewise purified Mer3 from insect cells as previously described [32], and demonstrated that both samples were free of detectable nucleic acid contamination as determined by the ratio of the absorbance at 260 nm and 280 nm in size exclusion chromatography (Supplementary Figure 1A-D). In a pull-down on magnetic streptavidin beads using C-terminally Strep-tagged Mer3 we found that Mer3 was able to interact directly with RPA (Figure 1E). This suggests that under these conditions, interaction between Mer3 and RPA can be established directly, and without ssDNA.

Mer3 is widely conserved across eukaryotes[36], and we therefore asked whether human HFM1 (hHFM1) could bind to human RPA (hRPA). hHFM1 has a 24% sequence identity and a 38% sequence similarity to yeast Mer3. In yeast two-hybrid assays, we detected an apparent interaction between HFM1 and human RFA1 (Figure 1F). Likewise, we found that mouse HFM1 (mHFM1) binds to human RFA1 in Y2H (Supplementary Figure 2). To exclude the possibility of an indirectly mediated interaction we also produced recombinant hHFM1 and recombinant human RPA (Supplementary Figure 1F and G). In a pulldown experiment on streptavidin beads we noticed an enrichment of MBP-hHFM1 in the presence of RPA versus the control, indeed pointing to a direct interaction between these proteins (Figure 1G).

We initially characterised the DNA binding properties of the Mer3-RPA complex, using both biotinylated 90-mer ssDNA and biotinylated reconstituted D-loops to explore DNA binding. We found that on D-loop DNA we could obtain a near stoichiometric complex with Mer3 and RPA (Figure 1H), whereas with ssDNA we clearly obtained more RPA (Supplementary Figure 3A). We also utilised EMSAs (Supplementary Figure 3B) which showed that Mer3 and RPA were able to form complexes on ssDNA at a slightly lower concentration of Mer3 when in the presence of 10 mM RPA. Taken together these data suggest that Mer3 and RPA do not clearly compete for DNA binding, and that there may be more of a cooperative assembly on DNA substrates.

To gain understanding of the structural organisation of the Mer3-RPA complex we analysed the Mer3-RPA complex by chemical cross-linking, with the DSBU cross-linker, coupled to mass-spectrometry (XL-MS) [37]. Despite obtaining many inter-polypeptide cross-links, we only found cross-links between Mer3 and Rfa1 and not between Mer3 and Rfa2 and Rfa3 (Figure 1I). The cross-linking data also indicated that the stoichiometry of the Mer3-RPA complex is higher than 1:1:1:1. We found multiple self-crosslinks (for Mer3, Rfa1 and Rfa3, Supplementary Figure 4), where a peptide has cross-linked to itself. This is not entirely surprising since we have previously shown that Mer3 can form dimers [32], though RPA is typically a 1:1:1 heterotrimer [6]. Regarding a potential RPA interaction site on Mer3, we note a particular accumulation of cross-links between the C-terminal Ig-like domain of Mer3 and the N-OB and DBD-A domains of Rfa1 (Figure 1I). This arrangement suggests that Rfa1 provides the Mer3 binding site.

### Mer3 interacts directly with the N-OB domain of Rfa1

To gain further insights into the potential nature of the interaction between Mer3 and RPA we utilised AlphaFold2 (AF2), run in multimer mode [38,39]. We did attempt to use the XL-MS data to validate the models, but this was challenged by the observation that many cross-links were not compatible with the model. We reasoned that this is because of the aforementioned higher-order stoichiometry (>1:1:1:1), which makes unambiguous interpretation of cross-linking data challenging. We also attempted to use the XL-MS dataset as restraints for AF2 modelling using AlphaLink [40,41], but the models were unchanged from the original prediction. The models we generated (Figure 2A and B) proposed one high-confidence interaction between Mer3 and RPA, which suggested that a C-terminal region of Mer3 (residues 992-1010) could bind to the N-OB domain of Rfa1 (Figure 2B). The N-OB domain of Rfa1 is the interaction site for various RPA interaction factors, including RAD2, MRE11, Ddc2 (ATRIP) and DNA2 which seem to share the same mode of interaction [42–44] (Supplementary Figure 5). In agreement with this idea, using yeast two-hybrid we observed that the N-terminal 140 amino acids of Rfa1, which contains the N-OB domain, is sufficient to facilitate an interaction with Mer3 (Figure 2C).

**Figure 2.**
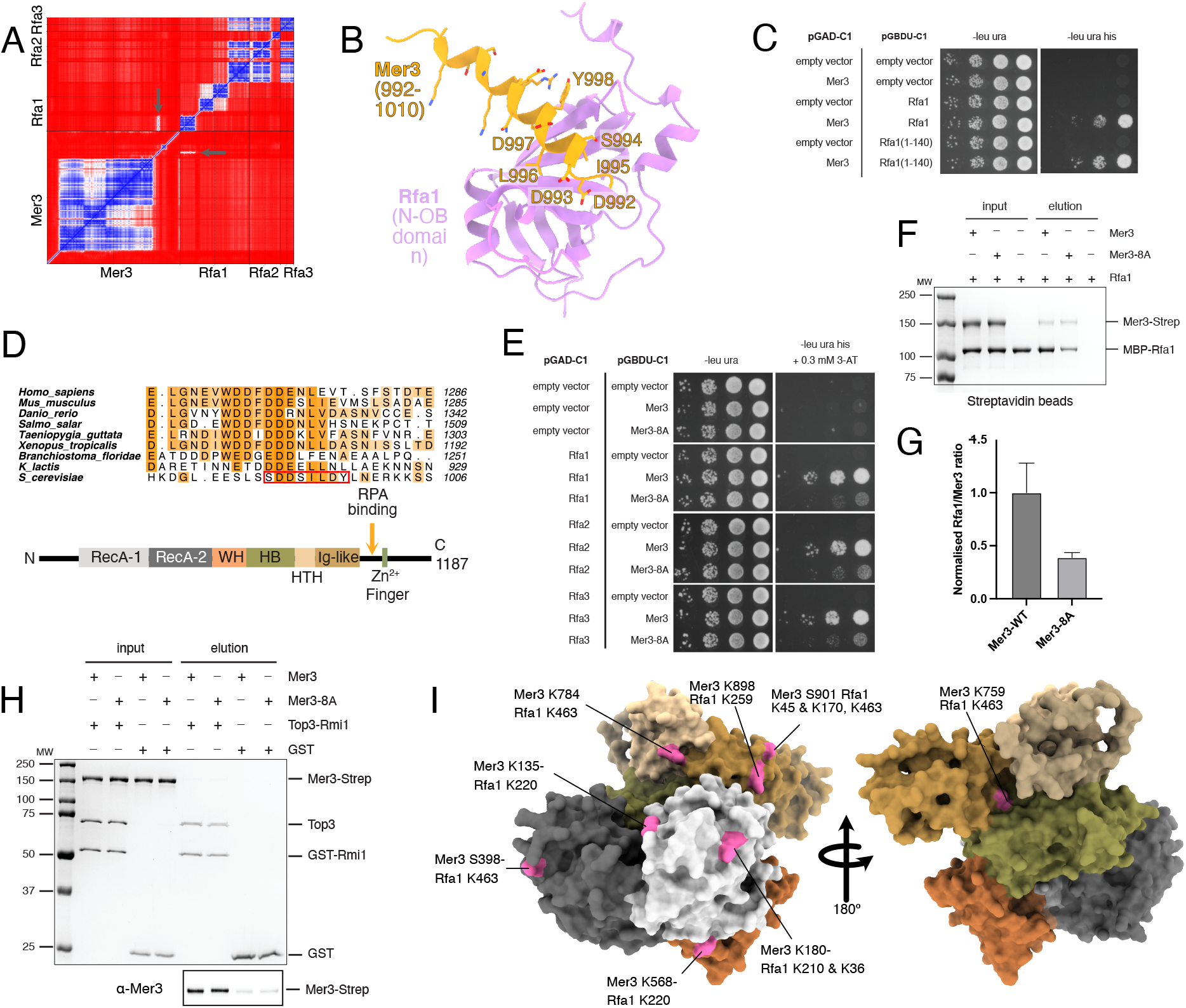
Mer3 binds to RPA via the N-OB domain. A) Predicted alignment error plot (PAE) from an AlphaFold2 run on Mer3-RPA. The plot suggests that the only interaction region of reasonable confidence that could be predicted is between the N-OB domain of Rfa1 and a C-terminal region of Mer3. B) Cartoon ribbon representation of the Mer3-RPA AlphaFold2 prediction. The C-terminal region of Mer3 (pale orange) is predicted to bind to a cleft in the N-OB domain of Rfa1 (pink). This type of interaction has been found in many other RPA binders (see Supplementary Figure 5). C) Yeast two-hybrid assay of Mer3 with full-length Rfa1 and the N-OB domain alone of Rfa1 (amino acids 1-140). D) Sequence alignment of interaction region. Alignment of Mer3/HFM1 sequences from selected species (mammals, other vertebrates, invertebrates, fungi), reveal that there is extensive conservation within the predicted binding region. The interaction region is shown in the domain cartoon of Mer3, coloured as in Figure 1A. E) Yeast two-hybrid assay of Mer3-8A mutant with RPA subunits. Mutation of the binding region to a poly-alanine sequence showed a substantial loss of association between Mer3 and all three of the RPA subunits. F) Pulldown of Strep-Mer3 and Strep-Mer3-8A against MBP-Rfa1 on Streptavidin beads. G) Quantification of the pulldown in F reveals a 2-fold loss of binding in Mer3-8A. Plotted values are means of three independent experiments. H) Pulldown of GST-Top3-Rmi1 against Mer3-WT and Mer3-8A shows that both the wild-type and the mutant bind equally well in the previously observed interaction [32]. I) Mapping of the Mer3-RPA XL-MS experiment onto the helicase core of Mer3 shows multiple cross-links between Mer3 and Rfa1, suggesting a second more widely dispersed interaction site on the catalytic region of Mer3.

Based on structural comparisons and multiple sequence alignment (Figure 2D) we identified 8 residues (DDSILDYL992-999) within Mer3 that are key to binding to the N-OB domain, both based on their position in the model and on their conservation (Figure 2D). We mutated these 8 residues to alanine (from hereon Mer3-8A) and tested whether an interaction between Mer3 and RPA could still be detected. In yeast two-hybrid experiments, we found that there was a noticeable, but not complete, loss of interaction between Mer3 and all three subunits of RPA (Figure 2E). Given that all three subunits of RPA are expressed at high levels in vegetative yeast, it still seems likely that the interaction between Mer3 and RPA is mediated primarily through Rfa1. To test this hypothesis we generated MBP-tagged Rfa1 alone and assayed its interaction with Mer3-WT and Mer3-8A in a pulldown experiment (Figure 2F). Under these conditions we could observe a robust interaction between Mer3-WT and MBP-Rfa1, which was reduced by at least 2-fold with Mer3-8A (Figure 2G). We attempted to obtain quantitative data, comparing the interaction between Mer3-WT and Mer3-8A with MBP-Rfa1. In a microscale thermophoresis experiment we found an approximate Kd of 1.31 μM +/-0.85 for Mer3-WT and 3.49 μM +/-1.03 for Mer3-8A, consistent with a >2-fold loss in binding affinity, though these results are complicated by elevated experimental noise (Supplementary Figure 6). For human HFM1, we made C-terminal truncations where the putative region was removed, and saw a total loss of interaction with hRFA1 (Figure 1F). We therefore propose that the mechanism of interaction is conserved. Curiously, although in yeast the RPA binding region is positioned N-terminal to the Zn^2+^ finger within Mer3, in mammals it lies C-terminal to the Zn^2+^ finger of HFM1. We carried out co-IPs on RPA in the Mer3-WT and Mer3-8A backgrounds, for which we observed no difference (Supplementary Figure 7A and B). While this was initially surprising, we surmised that this is presumably due to both the remaining lower affinity interaction between Mer3 and Rfa1, as seen above (Figure 2D-F), and to the presence of ssDNA in the pulldowns.

To ascertain the identified mutant was a separation-of-function mutant, and that other functions of Mer3 were not perturbed, we tested the ability of Mer3-8A to bind to other known partners of Mer3. In an experiment with the GST-Rmi1/Top3 complex, Mer3-8A was pulled down as effectively as Mer3-WT (Figure 2H). In yeast two-hybrid, we still observed an interaction between Mer3-8A and Dmc1 (Supplementary Figure 7C). Likewise, we found that our previously observed self-interaction for Mer3 was preserved in the Mer3-8A mutant [32] (Supplementary Figure 7D). We also examined DNA binding of this mutant. Mer3 has been shown to have a preference for D-loop DNA [30]. With reconstituted D-loops Mer3-8A and Mer3-WT both bound to the DNA substrate with equivalent apparent affinities (Supplementary Figure 7E). Taken together, we conclude that Mer3-8A specifically affects the interaction with Rfa1 of RPA, and that mutating these residues does not impact other known Mer3 interactions.

Are there structural changes that take place within Mer3 or RPA upon formation of the Mer3-RPA complex? We exploited our XL-MS approach to gain insights into changes in cross-linking patterns under different conditions. We carried out chemical cross-linking on our RPA complex alone (Supplementary Figure 8A), and compared this with the XL-MS data on the Mer3-RPA complex and our previous data from Mer3 in isolation [32]. The cross-linking maps of RPA alone and RPA with Mer3 are highly similar (Supplementary Figure 8A and B). However we do note that there are fewer cross-links between Rfa3 and the N-OB domain of Rfa1 in the presence of Mer3, which would be consistent with a scenario in which Mer3 engages the N-OB of Rfa1. Otherwise there is no detectable structural rearrangement of RPA when it binds to Mer3.

When we compared the cross-linking patterns within Mer3 in the presence or absence of RPA, we observed a greater concentration of short-range cross-links in the core of Mer3 (Supplementary Figure 8C and D). This indicates a likely structural rearrangement within Mer3, perhaps forming a less flexible or more globular structure. Consistently, although Mer3-8A showed a 3-fold loss of affinity for RPA, some interaction was retained in pulldowns, which might be through a direct, but low affinity, interaction with the helicase core of Mer3 (Supplementary Figure 8C and D).

In accordance with this idea, when we mapped the residues that cross-linked with RPA (Rfa1) onto the AlphaFold2 predicted structure of the helicase core of Mer3 we observed that the cross-links to Rfa1 are almost exclusively localised to the face of Mer3 where the two RecA-like domains are exposed to the solvent (Figure 2I). We thus propose that Mer3 contains two distinct interaction sites for RPA: One high-affinity and specific site in the C-terminal region of Mer3 (which is impacted in our Mer3-8A mutant), and a lower-affinity site distributed across the two RecA-like domains.

### RPA stimulates Mer3 processivity at low forces

RPA has been shown to stimulate the activity of several helicases through direct protein-protein interactions [21,45]. Similarly, Mer3 displays enhanced activity in the presence of either RPA or bacteria SSB; although, this stimulation has been attributed to the prevention of DNA by SSBs rather than to a direct effect on the helicase itself [29]. Based on this precedence, and supported by our evidence of direct contacts between Rfa1 and the helicase core of Mer3 that might alter its conformation, we investigated whether RPA could directly modulate Mer3 helicase activity. Bulk strand-separation assays revealed no apparent difference between Mer3 and Mer3-8A (Supplementary Figure 7F and G). To resolve helicase parameters of Mer3 proteins at the single-molecule level, we employed magnetic tweezers to monitor the behaviour of individual Mer3 molecules in real time.

A DNA hairpin construct containing a 1238 bp dsDNA region and a 45 nt ssDNA region (Supplementary Tables 4 and 5) as a loading site for Mer3 was tethered by its ends between a magnetic bead and a functionalized glass surface, respectively (Figure 3A) [46]. In the absence of proteins, applied forces above 15 pN confirmed that the hairpin opened abruptly, and closed below 15 pN (Supplementary Figure 9A). To ensure that unwinding signals reflected enzymatic activity rather than spontaneous hairpin opening, all measurements were carried out at forces below 15 pN. Under these conditions, the addition of 50 pM concentration of Mer3 and 1 mM ATP should turn out in processive unwinding of the dsDNA hairpin in the 3^*′*^-5^*′*^ direction, observed as an increase in bead-to-surface distance corresponding to the conversion of dsDNA into ssDNA (Figure 3B).

**Figure 3.**
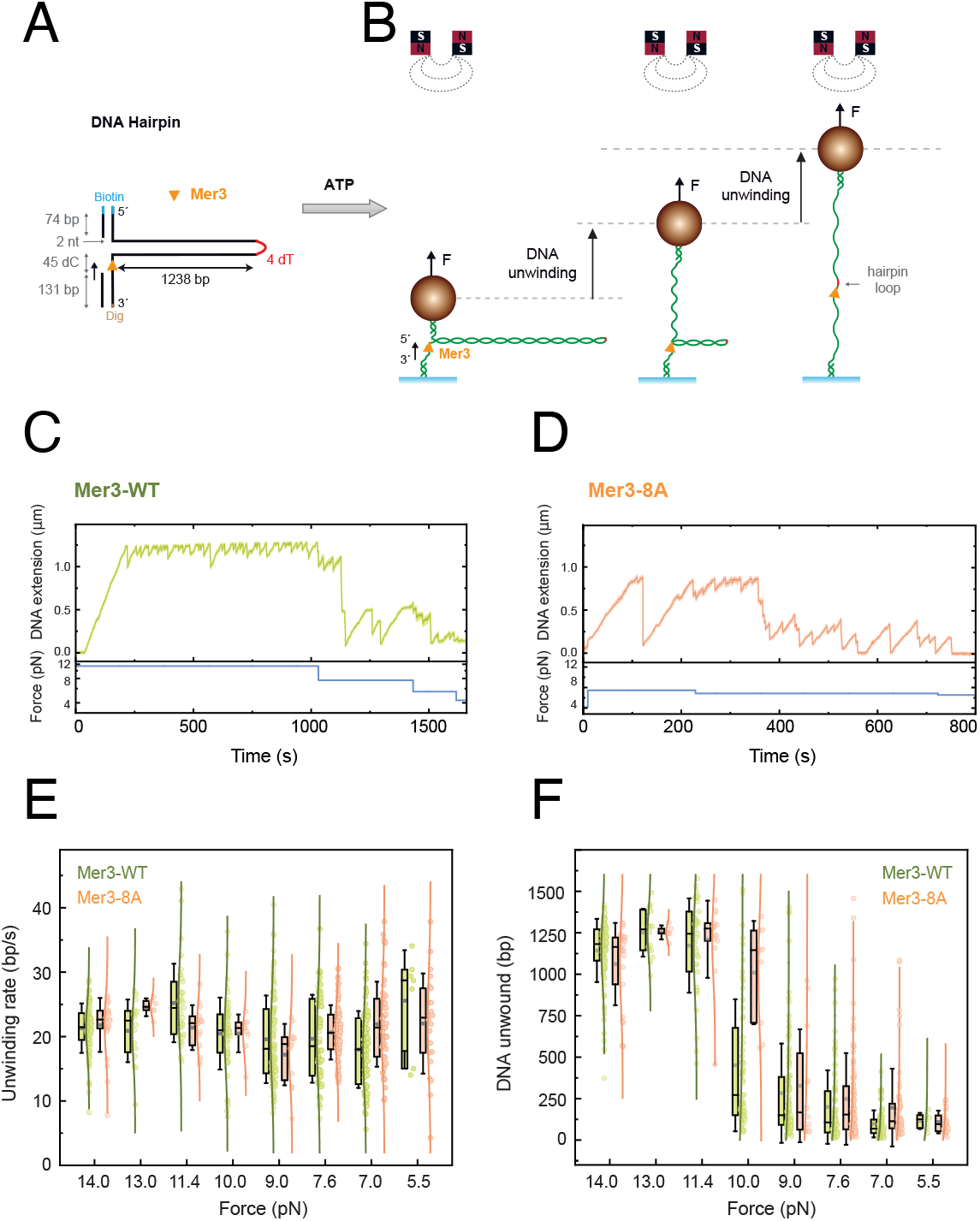
Single-molecule analysis of Mer3 helicase activity. A) Schematic of the DNA hairpin substrate used in magnetic tweezers experiments. B) Illustration of the single-molecule assay. Mer3 is loaded onto the single-stranded DNA region adjacent to the hairpin. Upon ATP addition, the helicase translocates in the 3–5 direction, unwinding the double-stranded region until it reaches the hairpin loop, resulting in complete opening of the hairpin. This assay enables precise quantification of unwinding rates and processivity under varying applied forces. C, D) Representative single-molecule activity traces measured at (C) 100 pM Mer3-WT and (D) 200 pM Mer3-8A, while the applied force was varied. E) Unwinding rates (bp/s) for Mer3-WT (green) and Mer3-8A (orange) at applied forces ranging from 14 to 5.5 pN. Both variants displayed a consistent unwinding rate of 22 bp/s independent of force. F) Length of DNA unwound (bp) for Mer3-WT (green) and Mer3-8A (orange) at the same range of forces. At higher forces, processivity reached 1200 bp, but it dropped substantially below 10 pN for both helicase variants.

**Figure 4.**
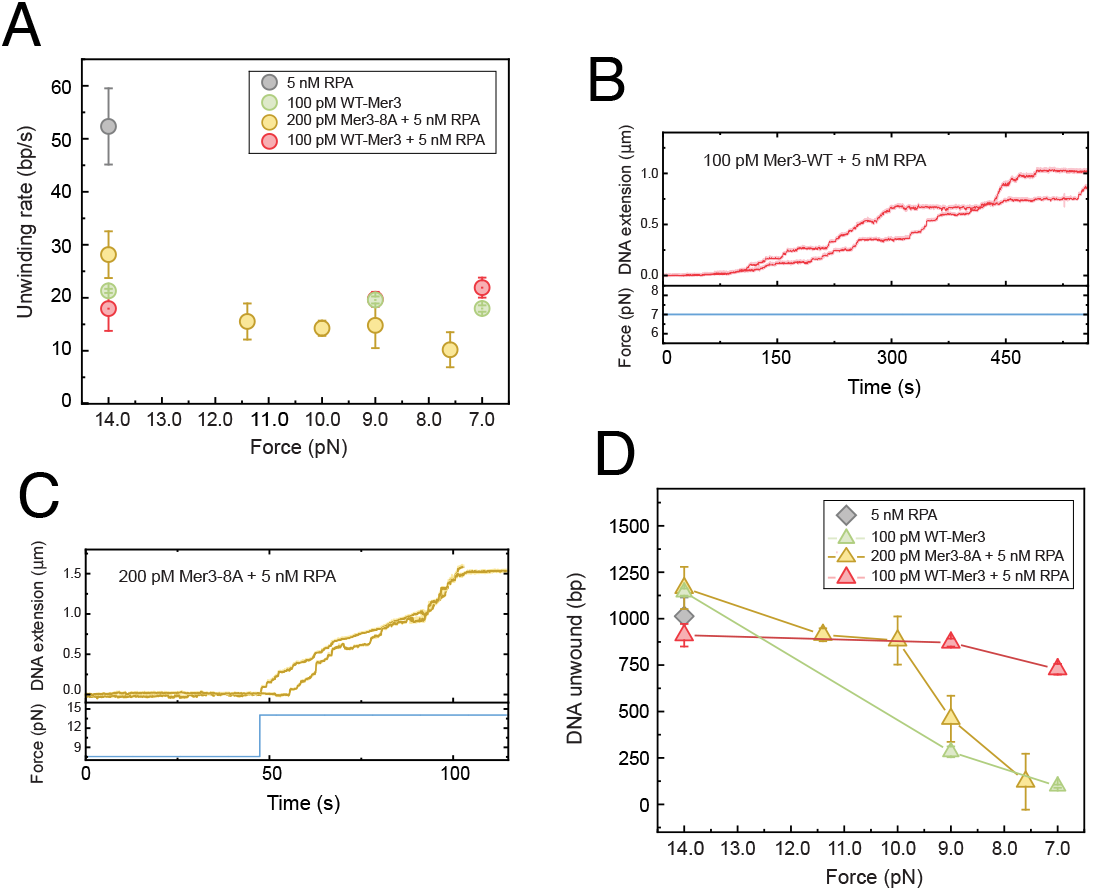
Mer3-RPA interaction is required for high processivity under low tension. A, B) Representative single-molecule activity traces for (A) 100 pM Mer3-WT with 5 nM RPA under 7 pN, and (B) 200 pM Mer3-8A with 5 nM RPA, with the applied forces varied from 7.6 pN to 14 pN. C) Effect of RPA on unwinding rate. Unwinding rates were measured for Mer3-WT (red circles) and Mer3-8A (yellow circles) in the presence of 5nM RPA across a range of applied forces (14 to 7pN). The presence of RPA had no detectable effect on the pause-free unwinding rate for either variant. D) Effect of RPA on processivity. Under the same conditions as in (C), processivity was assessed for Mer3-WT (red triangles) and Mer3-8A (yellow triangles). In the presence of RPA, Mer3-WT maintained high processivity across all forces tested, whereas Mer3-8A showed a pronounced decrease in processivity at forces below 10pN. For comparison, data for Mer3-WT alone (green) and RPA alone (grey) are also included in (C) and (D)

Representative single-molecule traces revealed unwinding activity by both Mer3-WT (Figure 3C) and Mer3-8A (Figure 3D) under applied forces below 15 pN. In both cases, DNA extension increased progressively until the hairpin was fully unwound. Following full hairpin unwinding, the helicase remained associated with the loop region (Figure 3A, red hairpin apex), often exhibiting a series of short, recurrent unwinding, translocation and slippage events resembling a sawtooth-like pattern.

To quantify helicase parameters, pause-free unwinding rates (see Methods) were measured across a range of force (from 14 pN to 5.5 pN). Both Mer3-WT and Mer3-8A exhibited similar and force-independent unwinding rates of 22 bp/s (Figure 3E). We also measured the total DNA length unwound prior to protein dissociation, used here as a proxy for processivity. Both helicase variants showed strong force dependence: at forces above 10 pN, processivity reached the full hairpin length (1200 bp), whereas at lower forces, processivity dropped sharply to 100 bp (Figure 3F). This suggests that applied force facilitates base-pair destabilization, effectively lowering the energy barrier for strand separation. As a result, helicase progression to unwind the duplex is easier and less likely to dissociate. Overall, these results indicate that the intrinsic helicase activity of Mer3 remained unaltered in the Mer3-8A mutant.

### RPA binding mutations cause cross-over defects

What effect might interfering with the interaction between Mer3 and RPA have on meiosis? To answer this, we mutated *MER3* to *mer3-8a* at the endogenous locus and monitored several aspects of meiotic recombination and progression through meiotic G2/prophase. We observed no effect on the timing of DNA replication (Figure 5A), and only a less than one hour delay of progression through meiosis I – based on expression of Cdc5 (Figure 5B and C) – in contrast to the severe defect seen for the *mer3*Δ mutant (Supplementary Figure 10A and B). We first examined the spore viability for *MER3, MER3-9xMyc, mer3-8A-9xMyc* and *mer3*Δ strains. We found that the presence of the C-terminal 9xMyc tag had no effect on spore viability, whereas the *mer3-8A-9xMyc* strain showed a moderately impaired, though significant reduction (p < 0.001), in spore viability of 87%, a nonetheless significantly healthier than the *mer3*Δ strains which showed a spore viability of 40% (Figure 5D) - consistent with previous reports of *mer3*Δ on spore viability [31]. Thus, on a global level, the *mer3-8A* allele showed a minor defect in meiotic behavior.

**Figure 5.**
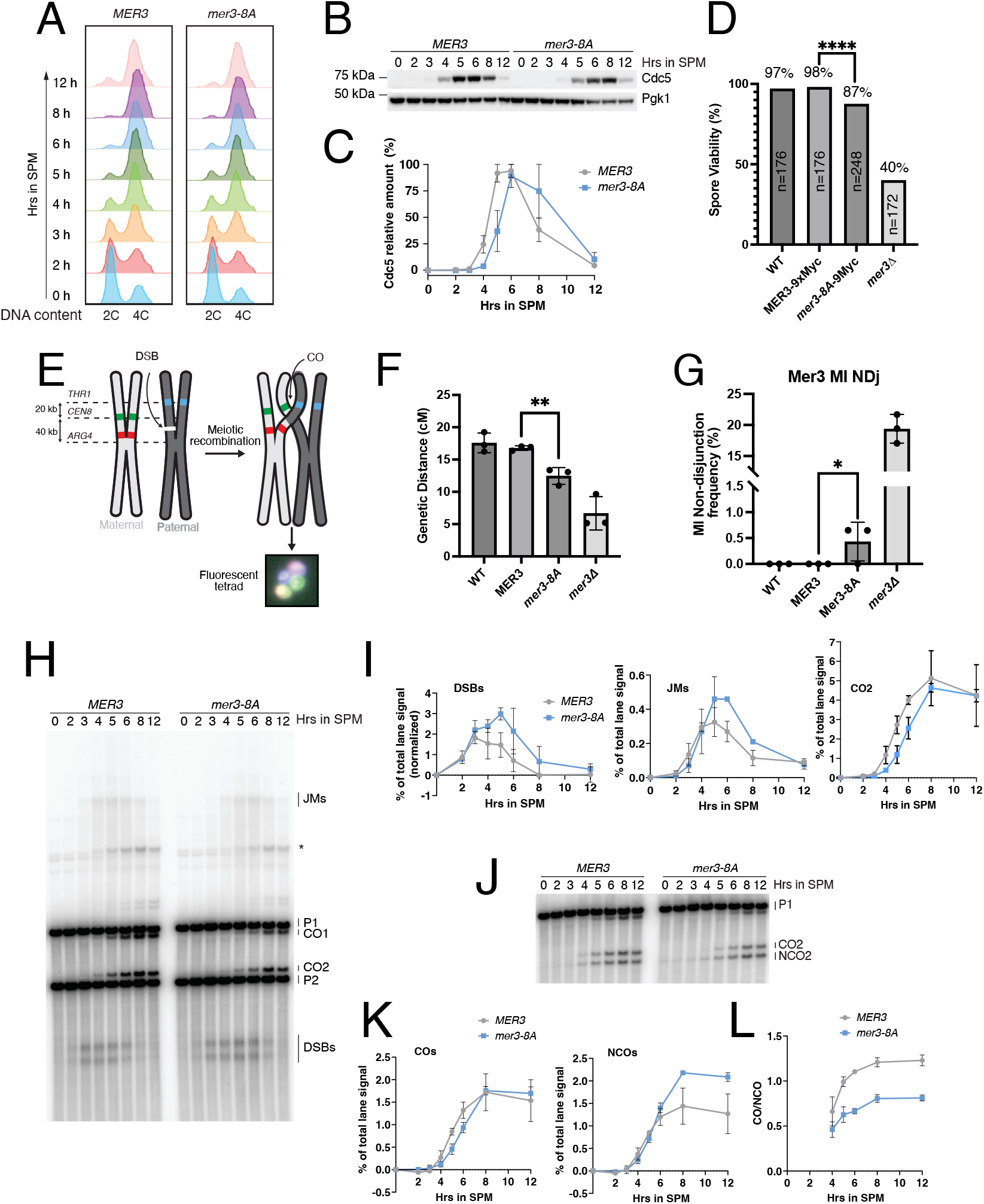
Mer3-8A mutant shows meiotic defects. A)) S. cerevisiae strains expressing MER3 or mer3-8A were synchronized in G0/G1 and released to undergo meiosis upon transfer to sporulation medium (SPM). Samples were collected at indicated time points to follow the progression of DNA replication. DNA content of the cells was analyzed by fluorescence-activated cell sorting. B) Cdc5 protein expression analysis from cells in (A). Protein samples were collected at indicated timepoints and analyzed by western blotting. C) Relative quantification of Cdc5 expression from (B) and a biological replicate shown in Supplementary Figure 10B, plotted as percentage of the peak Cdc5 signal. Plotted values are means of two independent experiments, error bars represent range. D) Spore viability. MER3 and MER3-9xMyc strains showed high spore viability (97% and 98% respectively). The mer3-8A-9xMyc strain showed a lower viability of 87.5%. The number of spores analysed by microdissection are indicated. Statistical significance between MER3-9xMyc and mer3-8A-9xMyc was determined by Fisher’s exact test, p=<0.0001. E) Cartoon of spore autonomous fluorescent assay, based on [49,50]. F) Genetic distances at the CEN8-ARG4 interval, based on three biological replicates. Statistical analysis of the difference between MER3 and mer3-8A was done with an unpaired t-test, resulting in a p-value of 0.005. G) MI non-disjunction events were also calculated from the spore autonomous fluorescence assay, using three biological replicates. Statistical significance between MER3 and mer3-8A determined by Fisher’s exact test (p=0.027). H) Physical analysis of recombination at HIS4-LEU2 in cells from A). DNA was psoralen-crosslinked and digested with XhoI, then analyzed by Southern blotting. JMs – joint molecules, P1 and P2 – parental DNA, CO1 and CO2 – reciprocal recombinants of P1 and P2, DSBs – double-strand breaks, asterisk represents ectopic crossing over. I) Double-strand break (DSB), joint molecule (JM) and crossover (CO2) quantification from H) and a biological replicate shown in Supplementary 10C. DSBs were plotted as percentage of total lane signal with subtracted background, joint molecules as percentage of total lane signal. Plotted values are means of two independent experiments, error bars represent range. J) Analysis of noncrossover and crossover formation at HIS4-LEU2. Psoralen-crosslinked DNA from H) was double-digested with XhoI and NgoMIV and analyzed by Southern blotting. P1 – parental DNA, CO2 – reciprocal recombinant of P1 and P2, NCO2 – noncrossover recombinant of P1 and P2. K) Crossover and noncrossover quantification from J) and a biological replicate shown in Supplementary Figure 10D, plotted as percentage of total lane signal. Plotted values are means of two independent experiments, error bars represent range. L) Ratio of crossovers to noncrossovers during meiotic progression calculated from data in K). Plotted values are means of two independent experiments, error bars represent range.

Since the interaction between Mer3 and RPA is likely neces-sary for Mer3 to process certain DNA intermediates during meiotic recombination, we asked whether the Mer3-8A mutant would lead to defective crossover formation. We utilised a previously described fluorescent-based assay to analyse genetic distance at the *CEN8-ARG4* interval and meiosis I non-disjunction frequency (Figure 5E) [49]. We found that the *mer3-8A* mutant strain had significantly reduced mean genetic distance between markers at the *CEN8* and *ARG4* loci of 12.5 cM, versus 17.5 cM for the wild-type and *MER3* strains (Figure 5F). As a reference the mer3 Δ strain had a genetic distance of 6.6 cM between the *CEN8* and *ARG4* markers (Figure 5F). Using the same assay we also observed a small fraction (0.5%) of non-disjunction events (MI NDJ), that is nonetheless significantly more than the zero NDJ events observed in the wild-type MER3 strains, and far smaller than in the mer3 Δ strain (20%) (Figure 5G). Thus, these data indicate that the *mer3-8a* allele disrupts normal meiotic crossover formation.

We next carried out a physical analysis of recombination at the well-studied *HIS4-LEU2* hotspot [51–55] using DNA extracted from yeast cultures at different time points through meiosis, that was subjected to a digestion with XhoI and Southern blotting (Figure 5H and biological replicate in Supplementary Figure 10E). We used Southern blotting to determine DSB formation, the accumulation of joint molecules (JMs), and crossover events at the locus. We observed increased DSB levels in *mer3-8A* mutant (though not as strong as in *mer3*Δ strains, Supplementary Figure 10C), and higher maximum levels of JMs, but no increase in COs at the *HIS4-LEU2* locus (Figure 5I). To reinforce these results, we double-digested the DNA extracted from this experiment with *XhoI* and *NgoMIV* to allow us to visualise COs and NCOs at the *HIS4-LEU2* locus [55] (Figure 5J and Supplementary Figure 10D). Analysis of two biological replicates clearly showed that while CO levels were the same in *MER3* and *mer3-8A* strains, there were more NCOs in the *mer3-8A* strains (Figure 5K): there was thus a clear change in NCO-CO ratios in *mer3-8A* with more DSBs processed as NCOs. Using 2D physical assays we sought to determine how joint molecules might be altered in *mer3-8A*. At both 4hr and 5hr time points we found an increase in both single-end invasions (SEIs) and double Holliday junctions (dHJs), though the ratio between these two DNA repair intermediates was apparently the same in both *MER3* and *mer3-8A* (Supplementary Figure 10 E-I).

To understand the meiotic phenotypes better, we sought to gain a genome-wide view of the behaviour of the *mer3-8A* mutant. We undertook calibrated ChIP-seq experiments for Mer3 in both wild-type and *mer3-8A* strains at two time points in meiosis. We constructed C-terminal His6Flag3 tagged strains of *MER3* and *mer3-8A* at the endogenous locus. Spore viability analysis of these strains confirmed the significant reduction in spore viability for the *mer3-8A-His6Flag3* strain (Supplementary Figure 11A). As a control, we analysed localisation of Mer3 in a *spo11*Δ strain, which is defective in double-strand break formation, and thus meiotic recombination. Mer3 is reported to have two broad localisation patterns:, one at Red1-enriched sites [58]. Red1 is a key component of the meiotic axis. The other localisation pattern of Mer3 corresponds to Spo11-dependent DSB hotspots [58]. We compared the localisation of our datasets across the genome (example from chromosome IX is shown in Figure 6A). We analysed Mer3 chromosomal locations at the strongest 500 DSB hotspots, the weakest 500 DSB hotspots and at mapped chromosomeaxis sites (e.g. Red1-associated sites). We found that in the *mer3-8A* strain there was weaker localisation of Mer3 at the DSB hotspots, but an apparently normal accumulation of Mer3 on the axis sites (Figure 6A-C, Supplementary Figure 11B-D, green traces)). In general Mer3 showed strongest association with DSB sites (Figure 6D, Supplementary Figure 11E), as previously shown [58]. Also as shown previously, the association of Mer3 with the axis sites required the formation of DSBs, since Mer3 did not associate with the axis sites in the *spo11*Δ background (Figure 6E, orange trace) [58]. This potentially reveals two kinds of “recruitment” of Mer3 to recombination sites: early (DSB-dependent) - visible strongly at the early stage of recombination (strongest association with axes) - preserved in the 8A mutant; and late (at DSB sites), corresponding to the “ZMM-stage” function of Mer3, with RPA - more intimately associated with the DSB hotspot sequence. These data were validated by qPCR at defined DSB hotspot and axis sites, at time points up to 8 hours in meiosis. We found that the Mer3-8A signal at DSB sites remained low throughout the meiotic time-course (Figure 6F and G and Supplementary Figure 11F).

**Figure 6.**
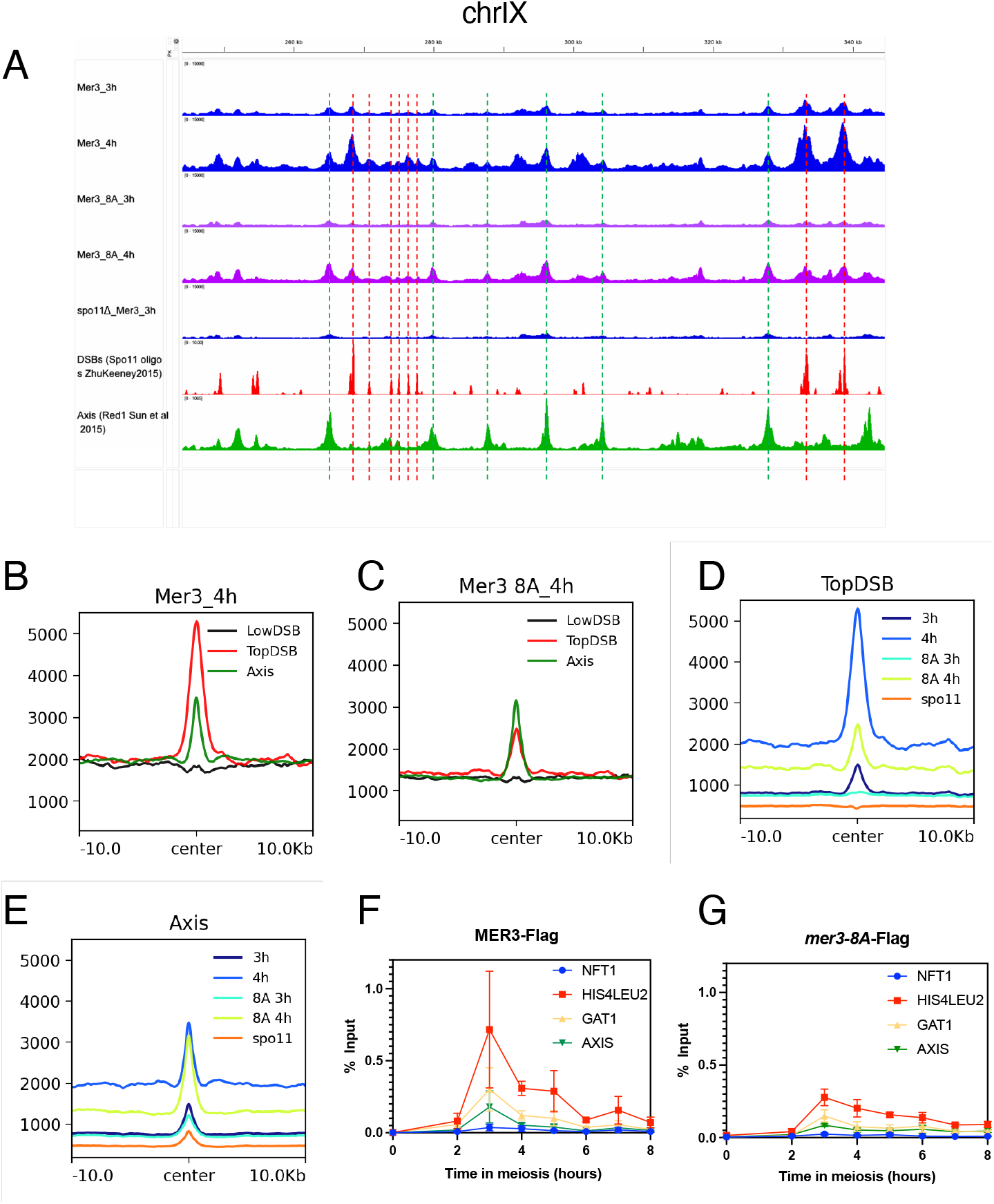
chIP-seq and chIP-qPCR reveal the deficient recruitment of Mer3-8A at DSB sites. A) ChIP-seq DNA binding for Mer3-His6Flag3 (blue) and Mer3-8A-His6Flag3(purple) at 3 hours and 4 hours in sporulation media as indicated. The DNA binding for Mer3-His6Flag3 in the *spo11*Δ background is shown as a control. DSB sites (Spo11 oligo signal) [52] are shown in red as well as axis association sites (Red1 ChIPseq, in green [53]). Red dotted lines are used to illustrate some DSB sites across the figure, likewise green dotted lines are used to indicate axis sites. B) Average Mer3-WT at 3 hours ChIP-seq signal of data shown in A at the indicated features. Alignments were performed on the Spo11 hotspots midpoints from[52] and Red1 peaks summits from [53]. C) Average Mer3-WT at 4 hours ChIP-seq signal of data shown in A at the indicated features. D) Comparison of the averages for Mer3-WT and Mer3-8A at 3 and 4 hours (and Mer3-WT at 3 hours in the *spo11*Δ background) for the strongest 500 Spo11 DSB hotspots. E) Average Mer3-8A at 3 hours ChIP-seq signal of data shown in A at the indicated features. F) Average Mer3-8A at 4 hours ChIP-seq signal of data shown in A at the indicated features. G) As E but with axis sites.

In summary, we identified a novel direct interaction between Mer3 and RPA, and found this interaction is conserved in other species. The binding of RPA and Mer3 stimulates Mer3 processivity on complex substrates and is necessary for normal crossover outcomes. We propose that the crossover defects arise due to a delayed processing of DNA repair intermediates, shown via both an accumulation of joint molecules, but also due to a shorter persistence of mutant Mer3 at the DSB sites.

## Discussion

Our data indicate that the 20-aa C-helix of Mer3 (residues 990-1010) binds the basic groove of Rfa1 N-OB, a site shared by Dna2, ATRIP and MRN and other RPA interactors [42,43]. Based on our work, we propose a two-step binding model: high-affinity Mer3 C-helix-mediated docking to Rfa1 positions RPA adjacent to the RecA-like lobes of Mer3; subsequent weaker, distributed interaction between Rfa1 and the helicase core could then act as a molecular clamp, reducing premature dissociation when the motor encounters secondary structures in ssDNA (Figure 7A). The unchanged unwinding rate but elevated processivity is consistent with such an allosteric reduction in the helicase off-rate rather than catalytic acceleration driven by such a Mer3-RPA constellation. We speculate that strong RPA coupling evolved to ensure extension of meiotic D-loops, through all types of DNA substrates, stabilising SEIs and facilitating their conversion to dHJs and subsequent COs. Furthermore, we speculate that binding of Mer3 to RPA could serve to sequester RPA from Sgs1, further antagonising Sgs1 activity, akin to the manner by which Mer3 binds to Top3 and Rmi1 [32].

**Figure 7.**
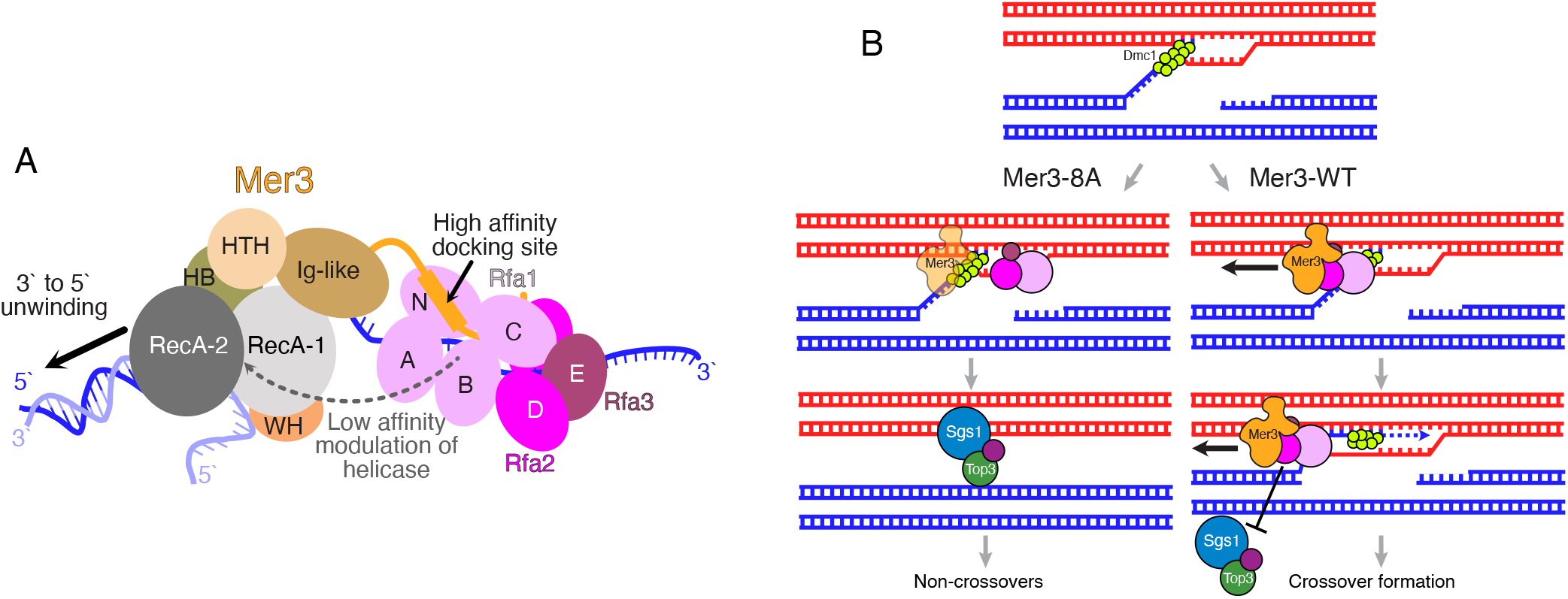
Model of the role of RPA binding to Mer3. A) Proposed domain organisation of Mer3 and RPA. Mer3 unwinds a DNA duplex in the 3 to 5 direction. RPA binds to exposed ssDNA via the DNA binding of Rfa1 DBD-A,B and C domains and the Rfa2 DBD-D domain. Mer3 and RPA dock via the binding of the Mer3 C-helix with the N-OB domain of Rfa1. The association between RPA and Mer3 allows further associations between the helicase core of Mer3 and Rfa1 to occur, which help to maintain the processivity of the helicase, particularly on complex DNA substrates. B) Mer3 is recruited to D-loops during meiotic recombination. In a Mer3-WT (right) the processivity of the Mer3-RPA complex allows the extension of D-loops and a longer Dmc1-mediated heteroduplex. This stabilises D-loops, favouring crossover formation. The stable association of Mer3 at the DSB sites also serves to protect these intermediates from disassembly by Sgs1-Top3-Rmi1 (STR). In the mer3-8A, Mer3-8A is only transiently associated with D-loops, since it cannot maintain a stable association with D-loops. This results in a shorter heteroduplex being formed, and a less stable intermediate. Furthermore the transient presence of Mer3-8A allows the STR complex to more readily disassemble these DNA repair intermediates, resulting in a higher number of NCO events.

In humans, biallelic HFM1 loss-of-function variants cause premature ovarian insufficiency and non-obstructive azoospermia, highlighting the clinical importance of this pathway [59]. Interesting, low-level HFM1 transcripts are detected in several tumour RNA-seq datasets (TCGA pan-cancer v33 [60]), raising the possibility that mis-expression of HFM1 outside of the germline could interfere with canonical RPA partner selection and DNA-repair fidelity under pathological conditions.

Taken together, a model for the role of the interaction between Mer3 and RPA and other factors is emerging. Mer3 is recruited early on in the recombination pathway to RPA-coated, Dmc1-mediated D-loops (Figure 7B). The physical association of Mer3 on Dmc1 mediated D-loops serves to stabilise these early recombination intermediates by protecting them from Sgs1/BLM [32]. The recombination intermediates are further stabilised by the helicase activity of Mer3, proposed to increase the length of the DNA heteroduplex formation [28]. The physical association between Mer3 and RPA serves to stimulate the processivity of Mer3 (Figure 7A), particularly on complex DNA substrates. Once the resected DNA is repaired by DNA polymerase delta, the interaction between Mer3 and Mlh1/Mlh2 serves to limit the length of gene conversion tracts [30]. Mer3 is likely also required for the recruitment of downstream ZMM proteins.

### Limitations and Future Directions

Stoichiometry of the Mer3–RPA complex remains ambiguous, as does the nature of the contact between RPA and the helicase core of Mer3. Thus, higher resolution structural studies, in the presence of a DNA substrate should be a goal for the future. A key next step will be to dissect how Mer3 processivity influences the recruitment of down-stream ZMM factors and the assembly of recombination nodules. Further dissection of mutant versions of Mer3, for example by comparing our *mer3-8a* allele with helicase-dead alleles and mutants that are defective for Mlh1 interactions should aid to illuminate this. Finally, testing whether human HFM1 requires RPA for enhanced processivity *in vitro*, and whether infertility-linked HFM1 variants disrupt this interface has the potential to connect the mechanistic insights gained from this current work to human disease.

## Materials and Methods

### Plasmids

Yeast gene ORFs were PCR-amplified from SK1 strain genomic DNA. Yeast MER3 and human HFM1 were amplified from codon-optimized gene blocks (IDT). Plasmids used for protein expression were cloned as previously described [57]. The mutations were introduced using gene-specific primers and subsequent Gibson assembly. Plasmids for yeast two-hybrid assay were prepared using the same protocol. PCR-amplified genes were cloned into yeast two-hybrid vectors pGAD-C1 or pGADCg (GAL4-activating domain), and pGBDU-C1 (GAL4-binding domain). The full list of plasmids used in this study is listed in the Supplementary Table 1.

### Protein purification

*Mer3* variants containing a C-terminal Twin-StrepII tag were produced in Hi5 insect cells. The cell pellet was resuspended in lysis buffer (20 mM HEPES pH 7.5, 300 mM NaCl, 5% glycerol, 0.1% NP-40, 5 mM betamercaptoethanol, AEBSF and Serva protease inhibitor cocktail) and sonicated. The lysate was clarified by ultracentrifugation at 35,000 rpm at 4°C for 1 h. Cleared lysate was applied on a 5 ml Strep-Tactin®XT column (IBA) followed by a first wash using 20 ml of H buffer (20 mM HEPES pH 7.5, 5% glycerol, 0.01% NP40, 1 mM β-mercaptoethanol) containing 500 mM NaCl and 1 mM ATP, and the second wash using 25 mL of H buffer containing 150 mM NaCl. The protein was eluted using 60 mL of H buffer containing 150 mM NaCl and 50 mM biotin. The fractions containing Mer3 protein were loaded onto a 5-mL HiTrap Heparin HP affinity column (Cytiva) pre-equilibrated in H buffer containing 150 mM NaCl and eluted with increasing salt gradient to 1 M NaCl. The peak fractions were concentrated on a 50 kDa MWCO Amicon concentrator and applied onto a Superdex 200 10/300 column (Cytiva) pre-equilibrated in SEC buffer (20 mM HEPES pH 7.5, 300 mM NaCl, 5% glycerol, 1 mM TCEP). The fractions containing Mer3 were concentrated on a 50 kDa MWCO Amicon concentrator and stored at −70°C in small aliquots. To obtain untagged and GFP-tagged Mer3 variants lacking Twin-StrepII tag, the concentrated fractions from Heparin column were mixed with 3C HRV protease in a molar ratio of 50:1 and incubated overnight at 4°C. Afterwards, the cleaved protein was loaded onto a Superdex 200 10/300 column (Cytiva) (pre-equilibrated in SEC buffer) with its outlet connected to a 5 mL GSTrap column (Cytiva) followed by 5 mL Strep-Tactin®XT column (IBA). The fractions containing Mer3 protein were concentrated on a 50 kDa MWCO Amicon concentrator and stored at −70°C in small aliquots.

*RPA* complex was produced in E. coli strain C41 by co-expression of pCOLI-6xHis-Rfa1, pCDF-6xHis-Rfa2 and pRSF-6xHis-Rfa3 plasmids. Protein expression was induced by 0.5 mM IPTG at 37°C for 3 h in TB media supplemented with ampicillin (100 μg/ml), kanamycin (25 μg/ml) and spectinomycin (50 μg/ml). The cell pellet was resuspended in the lysis buffer (50 mM HEPES pH 7.5, 300 mM NaCl, 5% glycerol, 0.1% NP-40, 5 mM β-mercaptoethanol, AEBSF, and Serva protease inhibitor cocktail). Resuspended cells were lysed by sonication before clearance at 35 000 rpm at 4°C for 35 min. Cleared lysate was loaded on a 5 ml Talon column (Cytiva) followed by the first wash using 10 ml of H buffer (20 mM HEPES pH 7.5, 5% glycerol, 0.01% NP40, 1 mM β-mercaptoethanol) containing 500 mM NaCl and 1 mM ATP, and the second wash using 25 mL of H buffer containing 150 mM NaCl. The protein was eluted with a 50-ml gradient of 0–500 mM imidazole in H buffer containing 150 mM NaCl. The fractions containing RPA complex were further loaded onto a 5 ml HiTrap Heparin HP affinity column (Cytiva) pre-equilibrated in H buffer containing 150 mM NaCl and eluted with increasing salt gradient to 1 M NaCl. The peak fractions were concentrated on a 10 kDa MWCO Amicon concentrator and applied onto a Superose 6 10/300 column (Cytiva) pre-equilibrated in SEC buffer (20 mM HEPES pH 7.5, 300 mM NaCl, 5% glycerol, 1 mM TCEP). The fractions containing the RPA complex were concentrated on a 10 kDa MWCO Amicon concentrator and stored at −70°C in small aliquots. To obtain untagged RPA complex, the concentrated fractions from Heparin column were mixed with 3C HRV protease in a molar ratio of 50:1 and incubated overnight at 4°C. Afterwards, the cleaved protein was loaded onto a Superdex 200 10/300 column (Cytiva) (pre-equilibrated in SEC buffer) with its outlet connected to a 5 mL GSTrap column (Cytiva). The fractions containing RPA were concentrated on a 10 kDa MWCO Amicon concentrator and stored at −70°C in small aliquots.

*Rfa1* containing an N-terminal 6xHis-MBP tag was produced in E. coli strain C41 cells. Protein expression was induced by 0.5 mM IPTG at 37°C for 3 h in TB media supplemented with ampicillin (100 μg/ml). The cell pellet was resuspended in the lysis buffer (50 mM HEPES pH 7.5, 300 mM NaCl, 5% glycerol, 0.1% NP-40, 5 mM β-mercaptoethanol, AEBSF, Serva protease inhibitor cocktail and 1 mM PMSF). Resuspended cells were lysed by sonication before clearance at 35 000 rpm at 4°C for 35 min. Cleared lysate was loaded on a 5 ml MBPTrap column (Cytiva) pre-equilibrated in H buffer (20 mM HEPES pH 7.5, 5% glycerol, 0.01% NP40, 1 mM β-mercaptoethanol) containing 150 mM NaCl. The column was first washed using 10 ml of H buffer containing 500 mM NaCl and 1 mM ATP followed by the second wash using 25 mL of H buffer containing 150 mM NaCl. The Rfa1 protein was eluted with a 50-ml gradient of 0–20 mM maltose of H buffer containing 150 mM NaCl. The fractions containing eluted protein were loaded onto a 5 ml HiTrap Heparin HP affinity column (Cytiva) pre-equilibrated in H buffer containing 150 mM NaCl and eluted with an increasing salt gradient to 1 M NaCl. The peak fractions were concentrated on a 50 kDa MWCO Amicon concentrator and applied onto a Superdex 200 10/300 column (Cytiva) pre-equilibrated in SEC buffer (20 mM HEPES pH 7.5, 300 mM NaCl, 5% glycerol, 1 mM TCEP). The fractions containing Rfa1 protein were concentrated on a 50 kDa MWCO Amicon concentrator and stored at −70°C in small aliquots.

*Human HFM1* containing an N-terminal 6xHis-MBP tag was produced in Hi5 insect cells. The cell pellet was resuspended in lysis buffer (20 mM HEPES pH 7.5, 300 mM NaCl, 5% glycerol, 0.1% NP-40, 5 mM beta-mercaptoethanol, AEBSF and Serva protease inhibitor cocktail) and sonicated. The lysate was clarified by ultracentrifugation at 35,000 rpm at 4°C for 1 h. Cleared lysate was applied on a 5 ml MBPtrap column (Cytiva) followed by a wash using 25 ml of H buffer (20 mM HEPES pH 7.5, 5% glycerol, 0.01% NP40, 1 mM β-mercaptoethanol) containing 150 mM NaCl. The protein was eluted with a 50-ml gradient of 0–20 mM maltose of H buffer containing 150 mM NaCl. The fractions containing HFM1 protein were concentrated on a 100 kDa MWCO Amicon concentrator and applied onto a Superose 6 10/300 column (Cytiva) pre-equilibrated in SEC buffer (20 mM HEPES pH 7.5, 300 mM NaCl, 5% glycerol, 1 mM TCEP). The fractions containing HFM1 were concentrated on a 100 kDa MWCO Amicon concentrator and stored at −70°C in small aliquots. *Human RPA* complex was produced in E. coli strain C41 by co-expression of pCOLI-Twin-StrepII-RFA1, pCDF-6xHis-RFA2 and pRSF-6xHis-RFA3 plasmids. Protein expression was induced by 0.2 mM IPTG at 18°C overnight in TB media supplemented with ampicillin (100 μg/ml), kanamycin (25 μg/ml) and spectinomycin (50 μg/ml). The cell pellet was resuspended in the lysis buffer (50 mM HEPES pH 7.5, 300 mM NaCl, 5% glycerol, 0.1% NP-40, 5 mM β-mercaptoethanol, AEBSF, and Serva protease inhibitor cocktail). Resuspended cells were lysed by sonication before clearance at 35 000 rpm at 4°C for 40 min. Cleared lysate was loaded on a 5 ml StrepTactin column (IBA) followed by the first wash using 10 ml of H buffer (20 mM HEPES pH 7.5, 5% glycerol, 0.01% NP40, 1 mM β-mercaptoethanol) containing 500 mM NaCl and 1 mM ATP, and the second wash using 25 mL of H buffer containing 150 mM NaCl. The protein was eluted using 50 mL of H buffer containing 150 mM NaCl and 50 mM biotin. The fractions containing RPA complex were further loaded onto a 6 ml ResourceQ column (Cytiva) pre-equilibrated in H buffer containing 150 mM NaCl and eluted with increasing salt gradient to 1 M NaCl. The peak fractions were concentrated on a 10 kDa MWCO Pierce concentrator and applied onto a Superdex 200 10/300 column (Cytiva) pre-equilibrated in SEC buffer (20 mM HEPES pH 7.5, 300 mM NaCl, 5% glycerol, 1 mM TCEP). The fractions containing the RPA complex were concentrated on a 10 kDa MWCO Pierce concentrator and stored at −70°C in small aliquots. Top3-Rmi1 complex was purified as described previously [32].

### Cross-linking mass spectrometry (XL-MS)

For XL-MS analysis Strep-tagged Mer3 and untagged RPA proteins were diluted in 200 μl of XL-MS buffer (20 mM HEPES 7.5, 300 mM NaCl, 5% glycerol, 1 mM TCEP) to the final concentration of 3 μM. The samples were mixed with 3 μl of disuccinimidyl dibutyric urea (DSBU) (200 mM) and incubated for 30 min at 25°C. The reaction was stopped by adding 20 μl of 1 M Tris–HCl pH 8.0 and incubated for additional 30 min at 25°C. The crosslinked samples were precipitated by addition of 4×volumes of 100% cold acetone followed by overnight incubation at −20°C. Samples were analysed as previously described [37], making use of MeroX [58] for data analysis, which only assumes that one of the two cross-linked residues must be a primary amine. Each sample was measured twice in independent cross-linking experiments. For interaction network visualisation XVis software was used and for visualization of the crosslinks on the PDBmodel PyXlinkViewer [59] and XMAS [60] was used. Each time a different cut-off for the cross-linking credibility was selected depending on the quality of the cross-linking data. In all figures only cross-links that were measured in two independent datasets are shown.

### AlphaFold

AlphaFold models of Mer3 and RPA were generated using AlphaFold2 version 2.3.2 run in multimer mode at the Max Planck Computing and Data Facility in Garching. Typical approaches involved running 5 independent predictions, and 5 models per prediction. Custom scripts were used to generate PAE plots from the predictions. All visualisations of protein structures were created using ChimeraX [61].

### Pull-down assays

For pull-down between Mer3 and RPA, RPA complex (1.9 μM) was incubated with Strep-tagged Mer3 (1 μM) in the reaction buffer (20 mM HEPES pH 7.5, 5% glycerol, 125 mM NaCl, 1 mM TCEP, 0.1% Tween-20) for 20 min at 30°C in the thermomixer (950 rpm). Prewashed magnetic streptavidin beads (3 μl) were then added to the samples and the mixtures were incubated for 2 min at 30°C in the thermomixer (750 rpm). Beads were washed twice with 100 μl of the reaction buffer. The proteins were eluted by boiling in 30 μl 2× SDS Laemmli buffer. The samples were loaded onto 14% SDS-PAGE, stained by Der Blaue Jonas gel dye and also analysed by western blot. After semi-dry western blotting, the membrane was incubated with anti-RFA (Agrisera AS07-214, 1:5,000). Following the incubation with the secondary goat anti-rabbit IgG peroxidase conjugate antibody (Merck, 401353), the proteins were visualized by chemiluminescence detection (WesternBright ECL, Advansta).

For pull-down between Rfa1 and Mer3 variants, Rfa1 (1.2 μM) was incubated with Strep-tagged Mer3 variants (1.2 μM) in the reaction buffer (20 mM HEPES pH 7.5, 5% glycerol, 250 mM NaCl, 1 mM TCEP, 0.1% Tween-20) for 20 min at 30°C in the thermomixer (950 rpm). Pre-washed magnetic streptavidin beads (3 μl) were then added to the samples and the mixtures were incubated for 2 min at 30°C in the thermomixer (750 rpm). Beads were washed once with 150 μl of the reaction buffer. The proteins were eluted by boiling in 30 μl 2× SDS Laemmli buffer. The samples were loaded onto 10% SDS-PAGE and stained by Der Blaue Jonas gel dye. The gels were analyzed using ImageJ software. For pull-down between Top3-Rmi1 (containing GST-tagged Rmi1) and Mer3 variants, Top3-Rmi1 (1 μM) was incubated with Strep-tagged Mer3 variants (1.2 μM) in the reaction buffer (20 mM HEPES pH 7.5, 5% glycerol, 250 mM NaCl, 1 mM TCEP, 0.1% Tween-20) for 20 min at 30°C in the thermomixer (950 rpm). Pre-washed magnetic glutathione beads (1 μl) were then added to the samples and the mixtures were incubated for 1.5 min at 30°C in the thermomixer (750 rpm). Beads were washed once with 150 μl of the reaction buffer. The proteins were eluted by boiling in 30 μl 2× SDS Laemmli buffer. The samples were loaded onto 10% SDS-PAGE and stained by Der Blaue Jonas gel dye. The gels were analyzed using ImageJ software. For pull-down between RPA and Mer3 on biotinylated DNA substrate, pre-washed magnetic streptavidin beads (3 μl) were incubated with biotinylated DNA (50 nM) in the reaction buffer (20 mM HEPES pH 7.5, 5% glycerol, 200 mM NaCl, 1 mM TCEP, 0.1% Tween-20) for 15 min at 30°C in the thermomixer (750 rpm). The buffer containing unbound DNA was removed and RPA (1.1 μM) diluted in the reaction buffer was added to the reactions. After the incubation for 10 min at 30°C in the thermomixer (950 rpm), untagged Mer3 (1 μM) was added to the selected reactions and the samples were further incubated for 5 min at 30°C in the thermomixer (550 rpm). Beads were washed once with 150 μl of the reaction buffer. The proteins were eluted by boiling in 30 μl 2× SDS Laemmli buffer. The samples were loaded onto 10% SDS-PAGE and stained by Der Blaue Jonas gel dye. The gels were analyzed using ImageJ software. For pull-down between human HFM1 and RPA complex, HFM1 (110 nM) was incubated with RPA complex containing Strep-tagged RFA1 (1.1 μM) in the reaction buffer (20 mM HEPES pH 7.5, 5% glycerol, 125 mM NaCl, 1 mM TCEP, 0.1% Tween-20) for 20 min at 30°C in the thermomixer (750 rpm). Pre-washed magnetic streptavidin beads (3 μl) were then added to the samples and the mixtures were incubated for 2 min at 30°C in the thermomixer (750 rpm). Beads were washed once with 150 μl of the reaction buffer. The proteins were eluted by boiling in 30 μl 2× SDS Laemmli buffer. The samples were loaded onto 10% SDS-PAGE and stained by Der Blaue Jonas gel dye. For the western blot analysis, the samples were loaded onto a 10% or 14% SDS-polyacrylamide gel, respectively, and blotted to nitrocellulose membrane. Antibodies used were as follows: anti-6xHis (Proteintech, 66005-1-IG, 1:1,000), anti-MBP (New England Biolabs, E8032L, 1:1,000), anti-Strep-tag II (Abcam, ab76949, 1:1,000), goat anti-rabbit IgG peroxidase conjugate (Merck, 401353), goat anti-mouse IgG peroxidase conjugate (Merck, 401215). Signal was detected using WesternBright ECL (Advansta) and visualised by a ChemiDocMP (Bio-Rad Inc).

### DNA substrates

DNA substrates used in this study (fluorescently labelled and biotinylated D-loop, resp.) were prepared as described previously [32]. The sequences of all oligonucleotides are listed in Supplementary Table 3.

### Electrophoretic mobility shift assay

The DNA binding reactions (10 μL volume) were carried out in 25 mM HEPES pH 7.5, 0.1 μg/μL BSA, 60 mM NaCl, and 10 nM fluorescently labeled D-loop substrate (prepared as described in [32]). The reactions were started by addition of increasing amounts of Mer3 variants (18.7, 37.5, 75, and 150 nM) and incubated for 20 min at 30°C. After the addition of 2 μL of the gel loading buffer (60% glycerol, 10 mM Tris–HCl, pH 7.4, 60 mM EDTA, 0.15 % Orange G), the reaction mixtures were resolved in 0.9% agarose gel in 1x TAE buffer (40 mM Tris, 20 mM acetic acid, 1 mM EDTA, pH 7.5). The gels were scanned using Amersham Typhoon scanner (Cytiva) and quantified by ImageQuant TL software (Cytiva). In the DNA binding assays with RPA and Mer3, fluorescently labeled ssDNA (oWL985, 10 nM) was first incubated with or without RPA (10 nM) in 25 mM HEPES pH 7.5, 0.1 μg/μL BSA, 1 mM MgCl2, 60 mM NaCl for 5 min at 37°C followed by addition of increasing amounts of Mer3 (10, 20, 40, 80 nM). After the incubation at 30°C for 15 min, glutaraldehyde (0.05%) was added to each reaction followed by additional incubation for 5 min at 37°C. The reaction mixtures containing the ge loading buffer were resolved in 0.9% agarose gel in 1x TAE buffer and the gels were analysed as described above.

### Strand separation assay

The strand separation assay (10 μL volume) was carried out in 25 mM HEPES pH 7.5, 60 mM NaCl, 0.1 μg/μL BSA, 1 mM MgCl2, 1 mM ATP, 10 mM creatine phosphatase, 15 μg/ml creatine kinase, and 5 nM fluorescently labeled D-loop substrate (prepared as described previously [32]). The reactions were started by addition of increasing amounts of Mer3 variants (1.25, 2.5, 5, and 10 nM). After the incubation for 30 min at 30 °C the reactions were stopped with 0.5 mg/mL proteinase K and 0.1% SDS, and incubated for 5 min at 37°C. The samples were then mixed with 2 μL of the gel loading buffer (60% glycerol, 10 mM Tris–HCl, pH 7.4, 60 mM EDTA) and separated on 10% (w/v) native polyacrylamide gel in 1xTBE buffer at a constant voltage of 110 V for 1 h at 4°C. The gels were scanned using Amersham Typhoon scanner (Cytiva) and quantified by ImageQuant TL software (Cytiva).

### Microscale thermophoresis

Binding affinity analysis by microscale thermophoresis was performed at 25°C using the Monolith NT instrument (Nanotemper Technologies). All reactions (in triplicates) were done in the MST buffer (20 mM HEPES pH 7.5, 5% glycerol, 200 mM NaCl, 0.1% pluronic), and contained constant concentration of GFP-tagged Mer3 variant (75 nM) and increasing concentrations of His-tagged RPA complex. Data were analysed by the MO.Affinity Analysis software (NanoTemper Technologies).

### Yeast strains

All strains, except those used for Y2H analysis, were derived from S. cerevisiae SK1 strains YML4068 and YML4069 and their genotypes are listed in Supplementary Table 2. Strains containing C-terminally Myc9-tagged Mer3 at its endogenous locus were prepared by a standard PCR-based approach using plasmid pYM18 as described [62,63]. Point mutations in mer3-8A strains (DDSILDYL992-999AAAAAAAA) were introduced by mutagenic PCR and further confirmed by sequencing. Strains containing C-terminally His6-Flag3 tagged Mer3 at its endogenous locus are from [30]. To generate the mer3-8A-His6Flag3 allele, the mutation was introduced by mutagenic PCR followed by sequencing. The Mer3-8A-His6-Flag3 strain had 85.8% spore viability (104 tetrads) versus 95.4% in the Mer3-His6-Flag3 (p<0.00001, Fisher’s exact test). To generate strains for spore fluorescence assay and DNA physical assay, a stop codon was inserted into generated Myc9-tagged Mer3 strains using mutagenic PCR and plasmid pFA6-natNT2 thus replacing kanMX cassette by natNT2 cassette.

### Meiotic co-IPs

Meiotic time course was done as previously described [32]. 100 mL of meiotic cultures (at 5 hours into a meiotic time course) were harvested by spinning down at 3,000 rpm for 3 min followed by wash with 700 μL of cold H2O containing 1 mM PMSF. Cell pellets were resuspended in 700 μL of ice-cold co-IP buffer (50 mM Tris-HCl pH 7.5, 150 mM NaCl, 1% Nonidet P-40,1 mM EDTA pH 8.0, 1 mM PMSF, AEBSF, Serva protease cocktail and a Complete Mini EDTA-free cocktail of protease inhibitors which was freshly added) and glass beads. The cells were lysed using a FastPrep-24 disruptor (MP Biomedicals) (setting: 2x 40 sec cycles at speed 6.0). Lysates were cleared by 2 rounds of centrifugation for 10 min at 15,000 rpm and the supernatants were after each centrifugation step transferred to a clean microcentrifuge tube. 0.5 μL of antibody (anti-RFA; Agrisera AS07-214) was added to 300 μL of the lysate followed by 3 hours incubation at 4°C. As a negative control, 300 μL lysate was incubated without the antibody. Subsequently, 20 μL of buffer-washed Dynabeads Protein G (Thermo Fisher Scientific) was added and the samples were incubated overnight at 4°C. The next day, Dynabeads were washed four times with 500 μL of ice-cold IP buffer. For the final wash, beads were transferred to a new microcentrifuge tube and washed 500 μL of ice-cold IP buffer without Nonidet P-40. The beads were resuspended in 55 μL of 2x SDS Laemmli buffer and incubated for 5 min at 95°C. The samples were loaded onto a 8% SDS-polyacrylamide gel and blotted to nitrocellulose membrane. Antibodies used were as follows: anti-PGK1 (22C5D8, Thermo Fisher Scientific, 459250, 1:1,000), anti-RFA (Agrisera AS07-214, 1:5,000), anti-Myc (Abcam, ab1326, 1:5,000), goat anti-rabbit IgG peroxidase conjugate (Merck, 401353), goat anti-mouse IgG peroxidase conjugate (Merck, 401215). Signal was detected using WesternBright ECL (Advansta) and visualised by a ChemiDocMP (Bio-Rad Inc).

### Meiotic whole cell protein analysis

Meiotic time course was done as previously described [32]. 10 mL of meiotic cultures were collected at specific time points and harvested by spinning down at 3,000 rpm for 3 min. Cell pellets were resuspended in 200 μL of ice-cold co-IP buffer (50 mM Tris-HCl pH 7.5, 150 mM NaCl, 1% Nonidet P-40,1 mM EDTA pH 8.0, 1 mM PMSF, AEBSF, Serva protease cocktail and a Complete Mini EDTA-free cocktail of protease inhibitors which was freshly added) and glass beads. The cells were lysed using a FastPrep-24 disruptor (MP Biomedicals) (setting: 2x 40 sec cycles at speed 6.0). Lysates were cleared by 2 rounds of centrifugation for 10 min at 15,000 rpm and the supernatants were after each centrifugation step transferred to a clean microcentrifuge tube. 160 μL of cleared lysate was mixed with 300 μL of TCA lysis buffer (300 mM NaOH, 7.4% β-mercaptoethanol), incubated for 10 min on ice and followed by addition of 120 μL of 20% trichloroacetic acid. After 10 min incubation on ice, the samples were centrifuged for 5 min at 15,000 rpm. The pellets were washed with 1 mL of cold acetone followed by drying for 30 min at RT. The pellets were then resuspended in 100 μL of 2x SDS Laemmli buffer and boiled for 5 min at 95°C. The western blot analysis was done as described for meiotic co-IPs.

### Yeast two-hybrid assay

Yeast two-hybrid plasmids were co-transformed into the S. cerevisiae reporter strain (yWL365; a kind gift from Gerben Vader) and plated onto the selective medium lacking leucine and uracil. For drop assay, 2.5 μl from 10-fold serial dilutions of cell cultures with the initial optical density (OD600) of 0.5 were spotted onto -Leu/-Ura (control) and -Leu/-Ura/-His plates with or without 0.1-0.3 mM 3-aminotriazole. Cells were grown at 30°C for up to 2-4 days and imaged.

### Spore viability

Yeast strains (yWL429, 514, 527, 686) were streaked out on YPD plates and grown for 2 days at 30°C. Cells were then transferred to SPM-plate and grown for 4 days at 30°C. Small amounts of cells were resuspended in 20 μL of Zymolyase 100T (1 mg/ml) and incubated for 10 min at 37°C followed by an addition of 50 μL of sterile water. The cell suspension was transferred onto a YPD plate for a tetrad dissection using a SporePlay+ dissection microscope (Singer Instruments). At least 172 spores were dissected for each strain. Spore viability was calculated as a percentage of the total number of viable spores.

### Meiotic time-courses

The time-courses were performed as described previously in [64]. In short, diploid cells were streaked on YP-glycerol plates (2% peptone (m/V), 1% yeast extract (m/V), 2% glycerol (V/V), 2% agar (m/V)) to grow single colonies for 2 days at 30°C. Single colonies were picked and plated to grow as small patches on YPD plates (24h, 30°C). Over the next 48 hours, patches were expanded into a lawn of cells on a single YPD plate, and then onto four YPD plates (30°C). The cells from four plates were collected and inoculated into 3.5 L pre-sporulation medium (2% peptone, 1% yeast extract, 2% KAc) at OD600 0.3 in 10 L fermenter system [65]. Cultures were incubated at 25°C for 14 h. The cells were then centrifuged and washed with sporulation medium (SPM - 2% KAc), before inoculating into 1 L SPM at OD600 3.5-4.0 to initiate meiosis. Meiotic cultures were incubated in fermenters at 30°C, collecting samples at indicated time-points (with the moment of inoculation into SPM defined as t = 0 h).

### FACS analysis of DNA content

1 mL of cells was collected at indicated time-points, fixed in 70% ethanol and stored at 4°C. The cells were spun down and washed once with 50 mM Tris-HCl (pH 7.5), then treated with 400 μg/mL RNase A for 3-4 h, at 37°C. The cells were then washed with FACS buffer (200 mM Tris-HCl (pH 7.5), 211 mM NaCl, 78 mM MgCl2), and finally resuspended in FACS buffer with added 55 μg/mL propidium iodide. Samples were sonicated briefly. Cell suspension was diluted 10-20 times in 1 mL 50 mM Tris-HCl (pH 7.5) and analyzed using FACSCalibur cytometer (Becton Dickinson) running CellQuest software. Flow cytometry data was analyzed using FlowJo software.

### Western blot analysis (Cdc5 and Pgk1)

Analysis of protein from cell lysates was performed as described previously [66]. Briefly, 9 mL of cells was collected at indicated timepoints. Protein extracts were obtained by cell disruption us-ing glass beads in 10% trichloroacetic acid. Proteins were precipitated, then resuspended in 2x NuPAGE sample buffer, neutralizing the acid with 1 M Tris. Samples were run on NuPAGE 4-12% Bis-Tris gels using MOPS running buffer (Invitrogen). Proteins were transferred to a 0.45 μM PVDF membrane (Amersham). Following primary antibodies were used for immunoblotting: Cdc5 (1:2500, 4F10, MediMabs) and Pgk1 (1:10000, 22C5D8, Invitrogen). Goat anti-mouse immunoglobulins conjugated to HRP (1:10000, P0447 Dako) were used as a secondary antibody. Signal was detected with ChemiDoc MP Imaging System (Bio-Rad). Quantification was performed in Fiji.

### Physical analysis of recombination

Physical analysis of recombination was performed as described previously in [51,67,68]. In brief, 50-100 mL of cells was collected and resuspended in 50 mM EDTA, 50 mM Tris-HCl pH 8 with 0.1 mg/mL psoralen. DNA was crosslinked in UVP Crosslinker CL-3000L at 3600 mJ/cm2 (Analytik Jena), while keeping cells on ice and mixing regularly. After crosslinking, cells were centrifuged and frozen in liquid nitrogen, then stored at −80°C. Genomic DNA was extracted using phenol-chloroform extraction. For joint molecule analysis (one-dimensional and two-dimensional), DNA was digested with XhoI. For crossover and noncrossover analysis, DNA was double digested with XhoI and NgoMIV. DNA concentration was measured with Qubit dsDNA Broad Range kit and normalized. For one-dimensional analysis, 1.5-2 μg DNA was loaded on 0.6 % agarose gels and ran at 60 V for 21 h in 1x TBE. For two-dimensional analysis, 1.5-2 μg DNA was loaded on 0.4 % gels and ran first at 30 V for 21 h in 1x TBE, then in the second dimension at 160 V for 4 h in 1x TBE containing 0.5 μg/mL ethidium bromide. DNA was transferred to a GeneScreen Plus (Revvity) membrane by alkaline transfer. The membranes were hybridized with a radioactively labeled probe overnight and then exposed to a phosphor screen. The screen was imaged with an Amersham Typhoon scanner. Different recombination species on one-dimensional gels were quantified using ImageQuant software. Background subtraction was done by subtracting the signal at the time point 0 from all measurements. Additional background subtraction was performed for DSBs due to high background signal coming from sheared DNA, by subtracting background lane signal quantified from an area equal in size to that of the DSB signal. Recombination species on two-dimensional gels were quantified relative to total DNA (signal from all species) using Fiji. Background subtraction was done manually by subtracting the background signal quantified from an equivalent area for each species.

### Spore autonomous fluorescence recombination analysis

Spore autonomous fluorescence analysis was performed as previously described in [48], with modifications. Briefly, diploids containing were streaked on a YP-glycerol plate and grown for 48 hours at 30°C. Single colonies were picked and grown on YPD plates as small patches (24 h, 30°C), and then expanded into larger patches (24 h, 30°C). The cells were put on 2% KAc plates to sporulate for at least 48 h at 30°C. A loop full of sporulated cells was resuspended in 0.75 mL liquid 2% KAc and sonicated. Samples were imaged in four channels on DeltaVision Ultra epifluorescence microscope (GE Healthcare) equipped with a 60x/1.42 UP-lanXApo Oil objective, a sCMOS camera under control of Acquire Ultra user interface (version 1.2.3).

### ChIP-qPCR and calibrated ChIP-seq

For synchronous meiosis, cells were grown in SPS presporulation medium and transferred to 1% potassium acetate with vigorous shaking at 30°C as described [69]. For qPCR analysis, at each meiotic time point, 2×108 cells were processed as described in [70]. Quantitative PCR was performed from the immunoprecipitated DNA or the whole-cell extract DNA (input) using a QuantStudio 5 (Applied Biosystems, Thermo Scientific) and analysed as described [70]. Results were expressed as % of DNA in the total input present in the immunoprecipitated sample. Primers for GAT1, HIS4LEU2, AXIS and NFT1 loci have been described [54]. For calibrated ChIPseq, 1×109 cells were processed as described [71] except that, for spike-in normalization, 1×108 (10%) S. mikatae cells of a single meiotic culture, harvested at 4 h in meiosis and fixed using the same procedure as for S. cerevisiae, were added to each sample before processing. Each experiment was performed in two independent replicates. Purified DNA was subjected to library preparation using the Accel-NGS 1S Plus DNA Library Kit (1S Unique Dual indexing kit, Cat. No. 19096) and sequenced using an Illumina NovaSeq 6000 instrument, generating paired-end 100 base-pair (bp) reads. Reads were aligned to the SaccCer2 S. cerevisiae S288C genome [54], and to the S. mikatae genome assembly [72], removing duplicate reads, and reads matching to the rDNA and the mitochondrial genome, using Bowtie to generate Bam files. The aligned experimental reads from the two independent replicates were then combined using MergeSamFiles to generate a single Bam file per condition. Reads that aligned on the S. cerevisiae genome but not on S. mikatae were defined as the experimental reads. For defining the spike-in normalization factor, we then determined the number of reads that did not align on the S. cerevisiae genome but aligned to the S. mikatae genome assembly, generating the spike-in reads. Bam files were converted to bigwig format using deepTools bamCoverage, with a bin size of 10 bp and with a scaling factor calculated as the number of experimental reads divided by the number of spike-in reads. Peaks for Red1 ChIP-seq and Spo11 oligonucleotides were from [52,53], respectively.

### DNA hairpin for magnetic tweezers

The design of a DNA hairpin substrate consisting of a 1238 bp hairpin with a 4 nt loop (4 dTs), a 74 bp dsDNA fragment ligated to a 126 bp highly biotinylated dsDNA handle, and a 146 bp 3-digoxigenin labeled dsDNA tail was based on a previous published construct with some modifications [73]. The digoxigenin labeled dsDNA fragment is connected to the hairpin through a ssDNA dC45 region. A 1238 bp hairpin was fabricated based on a previous design[73] with some modifications. The DNA construct consists of a 1238 bp hairpin with a 4 nt loop (4 dTs), a 74 bp dsDNA fragment ligated to a 126 bp highly biotinylated dsDNA handle, and a 146 bp 3-digoxigenin labeled dsDNA tail. The digoxigenin labeled dsDNA fragment is connected to the hairpin through a 45 nt poly dCs ssDNA region. The dsDNA hairpin was obtained by PCR amplification with Phusion High-Fidelity DNA Polymerase (Thermo Scientific) using Lambda DNA (NEB) as template and oligonucleotides that include different BsaI restriction sites in each side of the PCR fragment (see Supplementary Table X for oligonucleotides) followed by purification (QIAGEN) as described previously [45]. This region was selected by running a homemade software that computes the GC content of a given sequence and selecting a running window of 100 bp. After digestion with BsaI restriction enzyme (NEB) followed by purification, we obtained a dsDNA fragment of 1204 bp with a homogeneous GC content and unique non-palindromic 5,-overhangs. This restriction site was selected to avoid unspecific products of ligation in the later steps. A fork structure was formed by two partially complementary oligonucleotides, the 359.5-Phosphate BsaI and the 386.Template hairpin 45dC (Supplementary Table X). The two oligonucleotides were annealed by heating at 95°C for 5 min and cooling down to 20°C at a −1°C min-1 rate in hybridization buffer (10 mM Tris-HCl pH 8.0, 1 mM EDTA, 200 mM NaCl, and 5 mM MgCl2). The final fork structure contained a 5,-cohesive end compatible with one of the ends of the PCR fragment. The oligonucleotide 250.Loop hairpin was equally self-annealed to create a short dsDNA hairpin with a cohesive end compatible with the other end of the PCR fragment. The fork structure and the short hairpin oligo (5X excess of each) were overnight ligated to either end of the 1204 bp-PCR fragment by using T4 DNA ligase (NEB). Next, the ligated DNA structure was annealed with 10X excess of the oligonucleotide 360.3_BsaI_anneal_359 that was complementary to the oligonucleotide 359.5-Phosphate BsaI, by heating at 95°C for 5 min and cooling down to 20°C at a −1°C min-1 rate in annealing buffer (10 mM Tris-HCl pH 7.5, and 1 mM MgCl2). After hybridization, a new BsaI 5,-cohesive end that was compatible with the short highly biotinylated dsDNA handle was created. A 2 nt-gap remained next to the hairpin structure. The dsDNA handle was prepared by PCR (see Supplementary Table X for oligonucleotides) including 200 μM final concentration of each dNTP (dGTP, dCTP, dATP), 140 μM dTTP and 66 μM Bio-16-dUTP (Roche) using the plasmid pSP73-JY0 as template[74] followed by digestion with the restriction enzyme BsaI. Then, the DNA structure already hybridized with the oligonucleotide 360.3_BsaI_anneal_359 was overnight ligated with 10X excess of the biotinylated dsDNA handle. The ligated DNA construct was gel extracted to remove the excess of oligos and handle, and purified (QIAGEN). Finally, the purified structure was annealed with 10X excess of the oligonucleotide 253.Primer_for_Dig that was partially complementary to the oligonucleotide 386.Template hairpin 45dC, by heating at 95°C for 1 min and cooling down from 80°C to 10°C at a −0.5°C 10 s-1 rate in annealing buffer (10 mM Tris-HCl pH 7.5, and 1 mM MgCl2). The digoxigenin labeling required to attach the DNA hairpin construct to a glass surface via anti-digoxigenin antibodies was then incorporated by filling in the overhangs by DNA polymerase (Klenow Fragment [3^*′*^→5{textprimeexo-], NEB) in the presence of dATP, dCTP and dUTP-digoxigenin (Roche) for 1 h at 37°C followed by heat inactivation for 20 min at 75°C. The completed DNA hairpin construct was ready to use in MT experiments without further purification. EDTA pH 8.0 to 1 mM final concentration was added to preserve. DNAs were never exposed to intercalating dyes or UV radiation during their production and were stored at 4°C.

## Acknowledgements

We thank Franziska Müller and Petra Janning (Max Planck Institute of Molecular Physiology, Dortmund, Germany) for the generation of XL-MS data. Thanks to Susanne Astrinidis, Rahmiye Kürkcü and Jennifer Jüngling in the JRW laboratory for assistance with cloning. We thank Gerben Vader and members of the JRW lab for comments on the manuscript. Work in the JRW lab was funded by the Max Planck Society and the DFG (WE 6513/2–1). Work in the VB lab was funded by Agence Nationale de la Recherche (ANR-21-CE44-0009), Fondation ARC and La Ligue Contre le Cancer grants. We thank the Institut Curie NGS platform, supported by grants from ANR-10-EQPX-03, ANR10-INBS-09-08 and from the Cancéropôle Ile-de-France. Work in the F.M.-H. laboratory was supported by grants PID2023-146255NB-I00, funded by MICIU/AEI/10.13039/501100011033 and FEDER, EU, and TEC-2024/TEC-158, funded by the Autonomous Region of Madrid and co-funded by the European Social Fund and the European Regional Development Fund. We thank Javier Mendia-Garcia for all his work in setting up the hairpin assay in the F.M.-H. laboratory and for many insightful discussions. The Matos Lab was supported by the Austrian Science Foundation FWF (SFB Meiosis - 8807-B) and the European Research Council ERC (101002629).

## Supplementary Data

**Supplementary table 1.** Plasmids used in this study.

| Plasmid ID | Description | Reference |
| --- | --- | --- |
| pWL1333 | pLIB-Mer3-Strep | This study |
| pWL2479 | pLIB-eGFP-Mer3-Strep | This study |
| pWL2219 | pLIB-Mer3(DDSILDYL992-999AAAAAAAAA)-Strep | This study |
| pWL2614 | pLIB-eGFP-Mer3(DDSILDYL992-999AAAAAAAAA)-Strep | This study |
| pWL1097 | pLIB-Mer3(K167A)-Strep | Altmannova et. al 2023 NAR |
| pWL520 | pCOLI-6xHis-Rfa1 | This study |
| pWL526 | pCDF-6xHis-Rfa2 | This study |
| pWL527 | pRSF-6xHis-Rfa3 | This study |
| pWL2346 | pCOLI-6xHis-MBP-Rfa1 | This study |
| pWL856 | pBIG1a-Top3/GST-Rmi1 | Altmannova et. al 2023 NAR |
| pWL1541 | pLIB-6xHis-MBP-HFM1 | This study |
| pWL2376 | pCOLI-Strep-RFA1 | This study |
| pWL2392 | pCDF-6xHis-RFA2 | This study |
| pWL2393 | pRSF-6xHis-RFA3 | This study |
| pWL1565 | pGAD-C1 | Altmannova et. al 2023 NAR |
| pWL1564 | pGBDU-C1 | Altmannova et. al 2023 NAR |
| pWL2152 | pGADCg | This study |
| pWL1700 | pGAD-C1-Mer3 | Altmannova et. al 2023 NAR |
| pWL1716 | pGBDU-C1-Mer3 | Altmannova et. al 2023 NAR |
| pWL2189 | pGBDU-C1-Mer3(DDSILDYL992-999AAAAAAAAA) | This study |
| pWL2207 | pGADCg-Mer3 | This study |
| pWL1663 | pGAD-C1-Rfa1 | This study |
| pWL1819 | pGAD-C1-Rfa2 | This study |
| pWL1971 | pGAD-C1-Rfa3 | This study |
| pWL1664 | pGBDU-C1-Rfa1 | This study |
| pWL1965 | pGBDU-C1-Rfa1(1-140) | This study |
| pWL2213 | pGADCg-Dmc1 | This study |
| pWL2215 | pGADCg-hHFM1 | This study |
| pWL2628 | pGADCg-hHFM1(1-1270) | This study |
| pWL2639 | pGAD-C1-mHFM1 | This study |
| pWL1839 | pGBDU-C1-hRFA1 | This study |
| pWL1823 | pGBDU-C1-hMLH1 | This study |
| pWL2605 | pGBDU-C1-mMLH1 | This study |

**Supplementary table 2.**
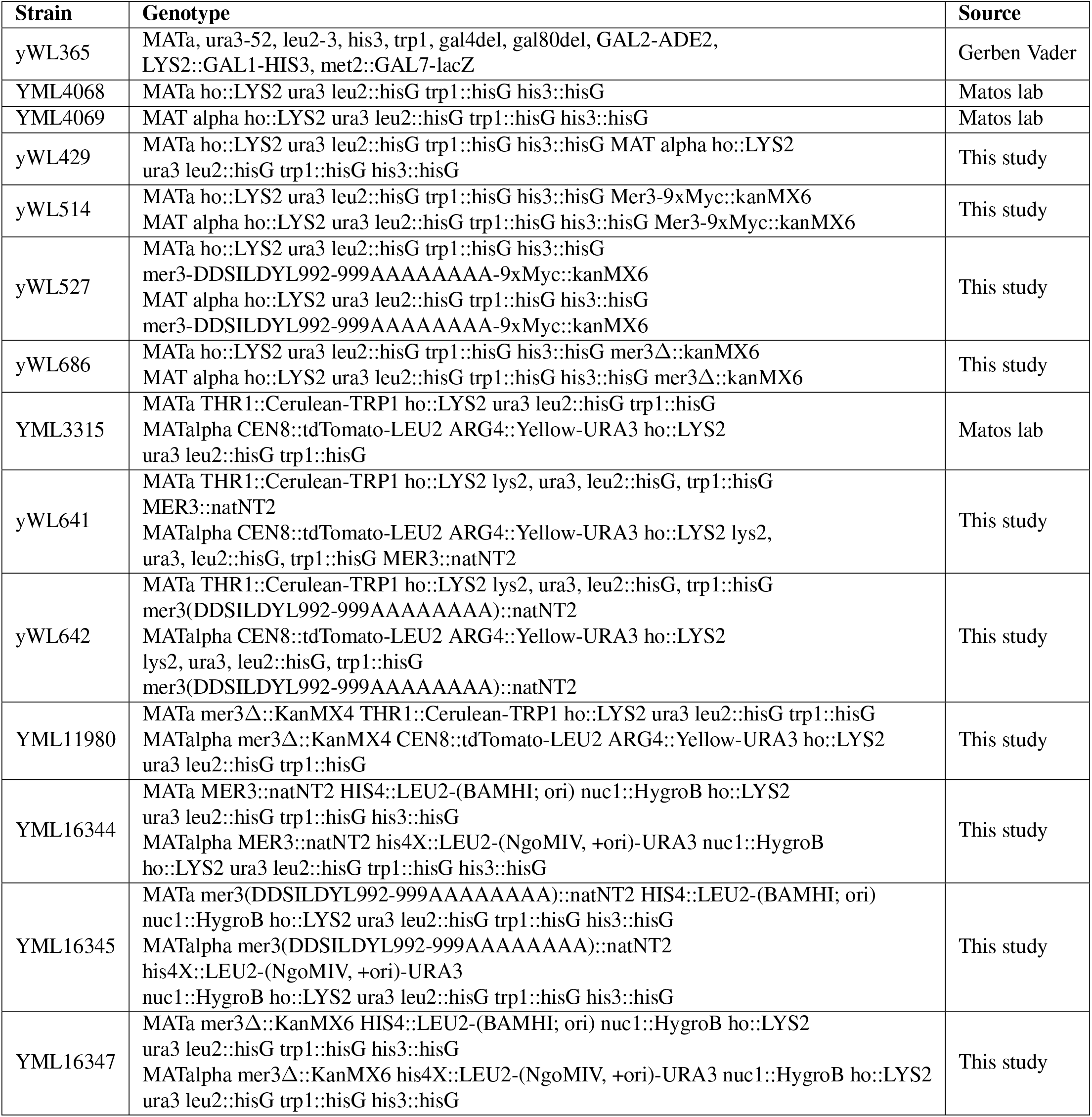
Yeast strains used in this study.

**Supplementary table 3.**
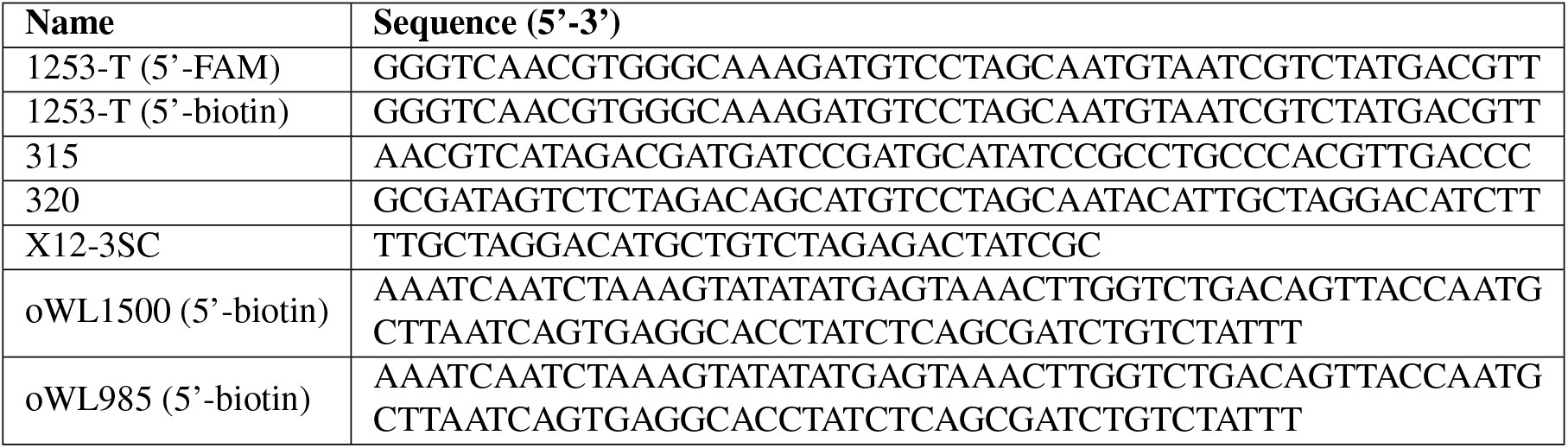
DNA oligonucleotides for EMSAs and Pulldowns.

**Supplementary table 4.**
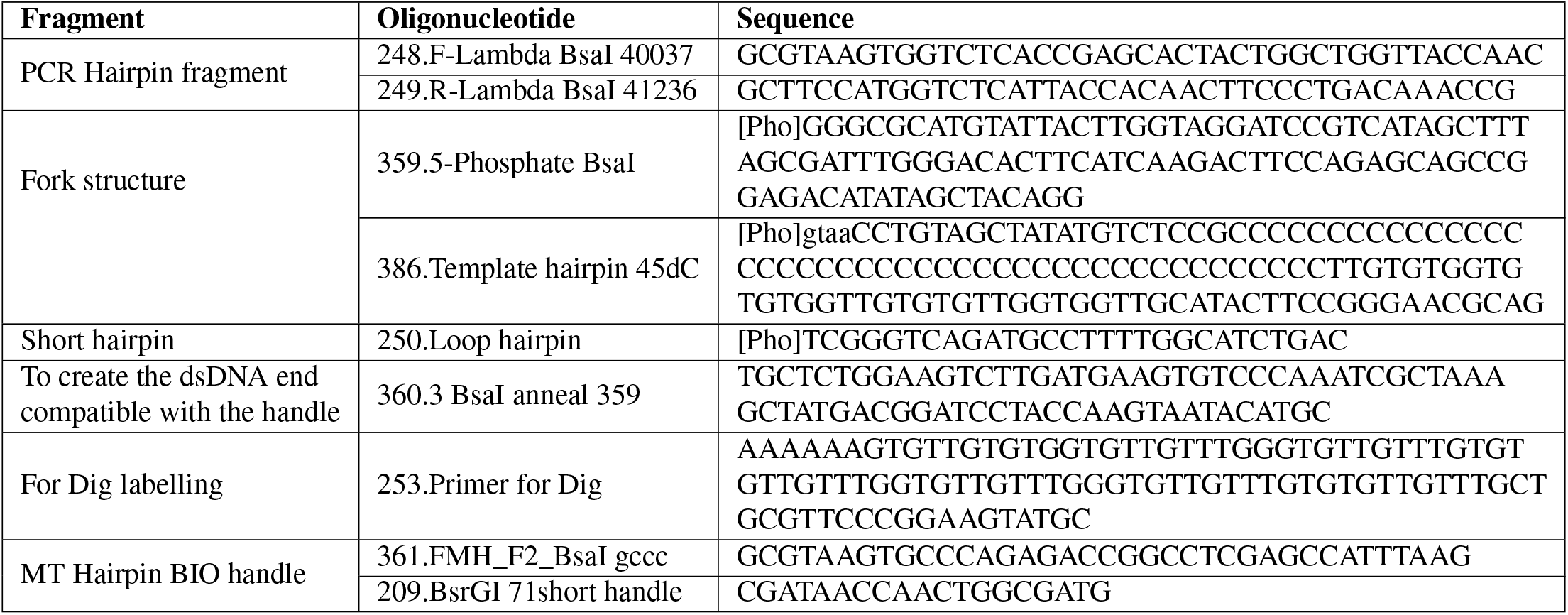
Sequences of oligonucleotides employed to fabricate the DNA Hairpin Substrate.

**Supplementary Table 5.**
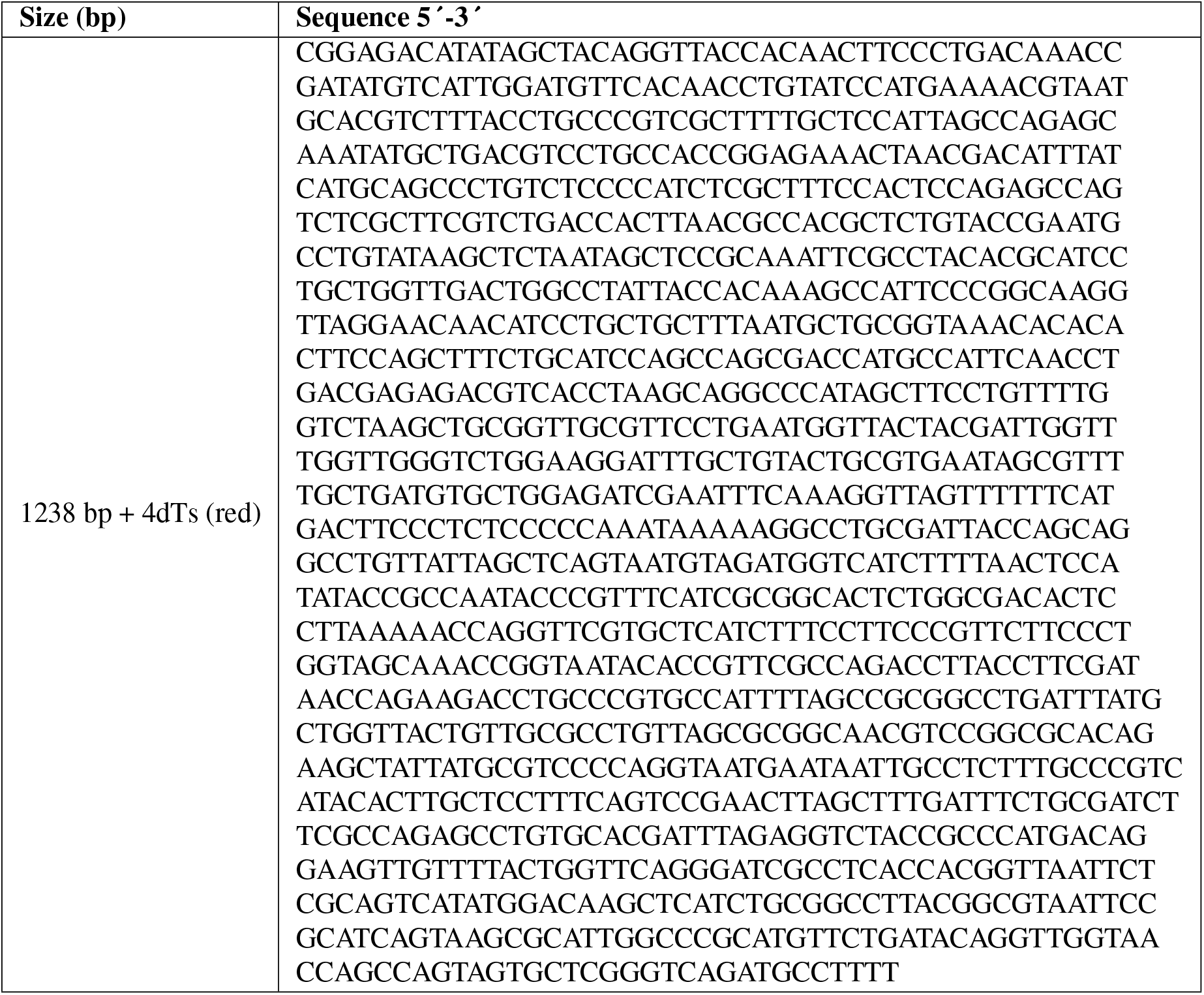
Sequence of the dsDNA central part of the final Hairpin Substrate.

**Supplementary Figure 1.**
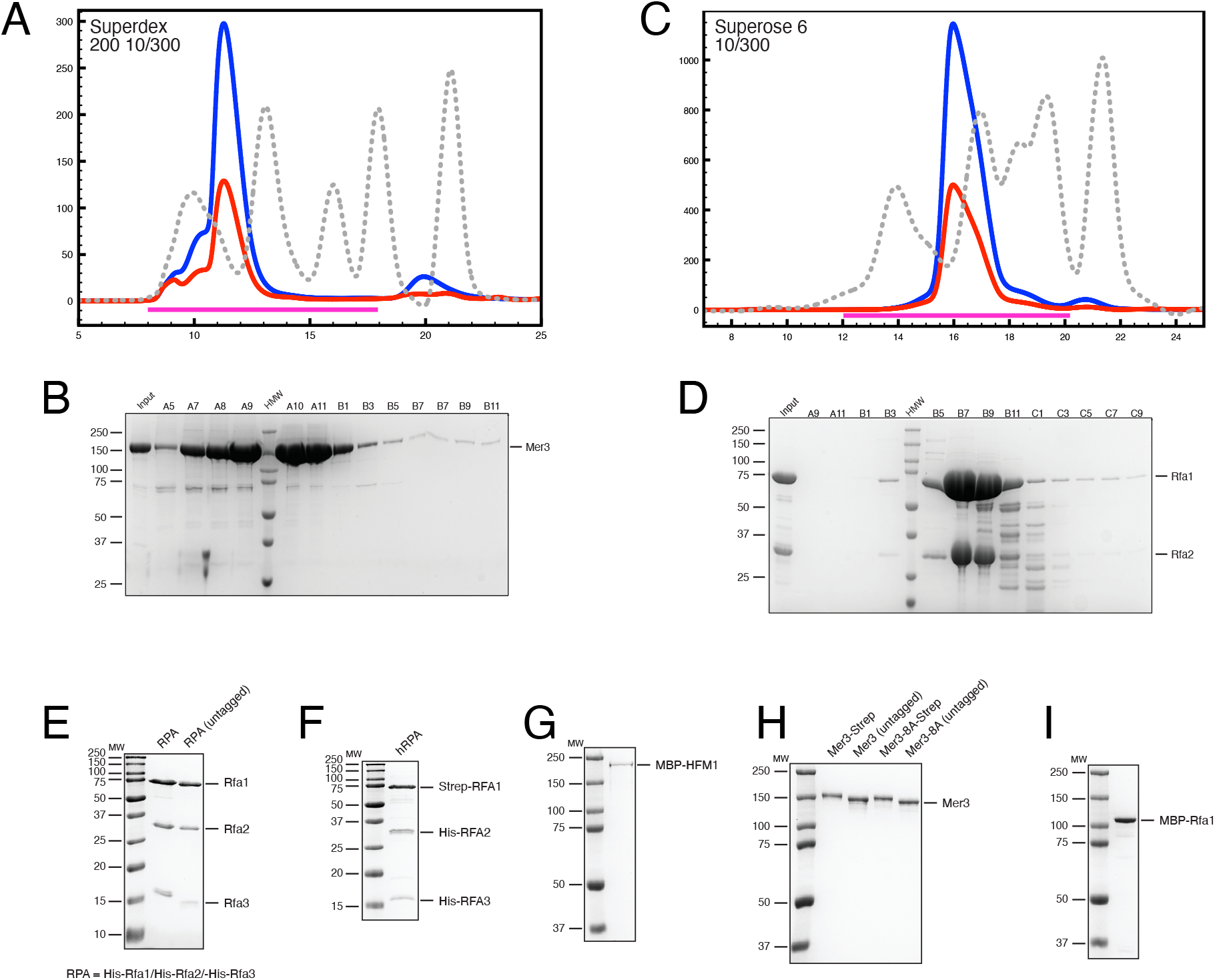
Purity of recombinant proteins and complexes. A) Size Exclusion Chromatography (SEC) profile of C-terminally Strep-tagged Mer3-WT run on a Superdex 200 10/300 column. The blue trace shows absorbance at 280 nm, the red at 260 nm. MW standards are included as a reference. Magenta bar indicates the region of fractions loaded in the gel shown in B). B) Coomassie stained SDS-PAGE of the SEC experiment shown in A). C) SEC profile of RPA run on a Superose 6 10/300 column. The blue trace shows absorbance at 280 nm, the red at 260 nm. MW standards are included as a reference. Magenta bar indicates the region of fractions loaded in the gel shown in B). D) Coomassie stained SDS-PAGE of the SEC experiment shown in C). E) Coomassie stained SDS-PAGE to demonstrate purity of tagged (left) and untagged (right) S. cerevisiae RPA complex. F) Coomassie stained SDS-PAGE to demonstrate purity of human RPA (hRPA) complex. G) Coomassie stained SDS-PAGE to demonstrate purity of N-terminally MBP-tagged human HFM1. H) Coomassie stained SDS-PAGE to demonstrate purity of different Mer3 constructs, as indicated. I) Coomassie stained SDS-PAGE to demonstrate purity of N-terminally MBP-tagged S. cerevisiae Rfa1.

**Supplementary Figure 2.**
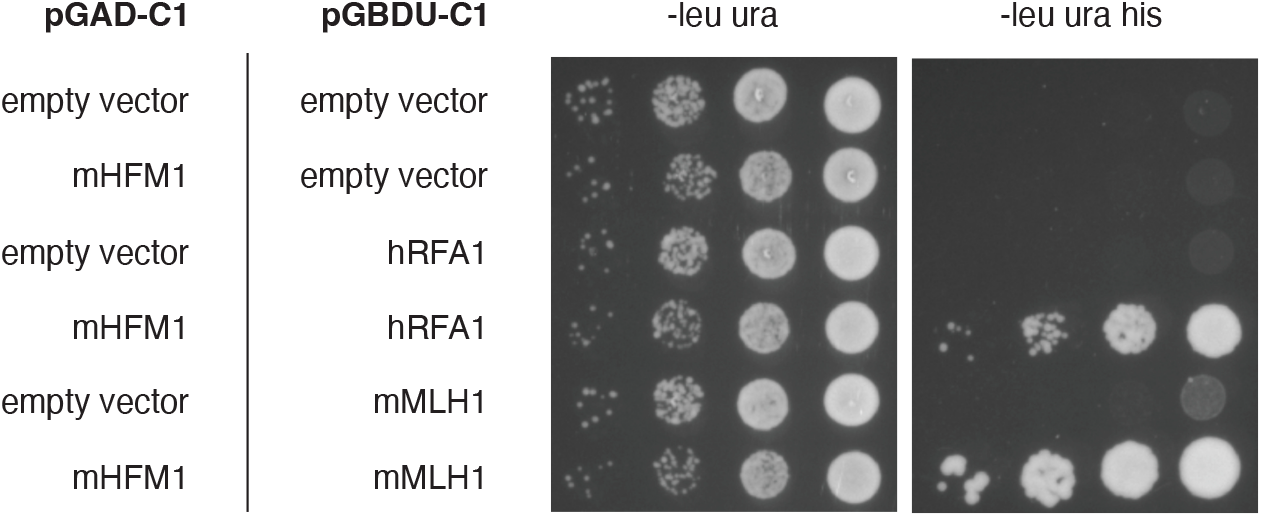
Mouse HFM1 interacts with human RFA1 in Y2H. mMLH1 was used as a positive control.

**Supplementary Figure 3.**
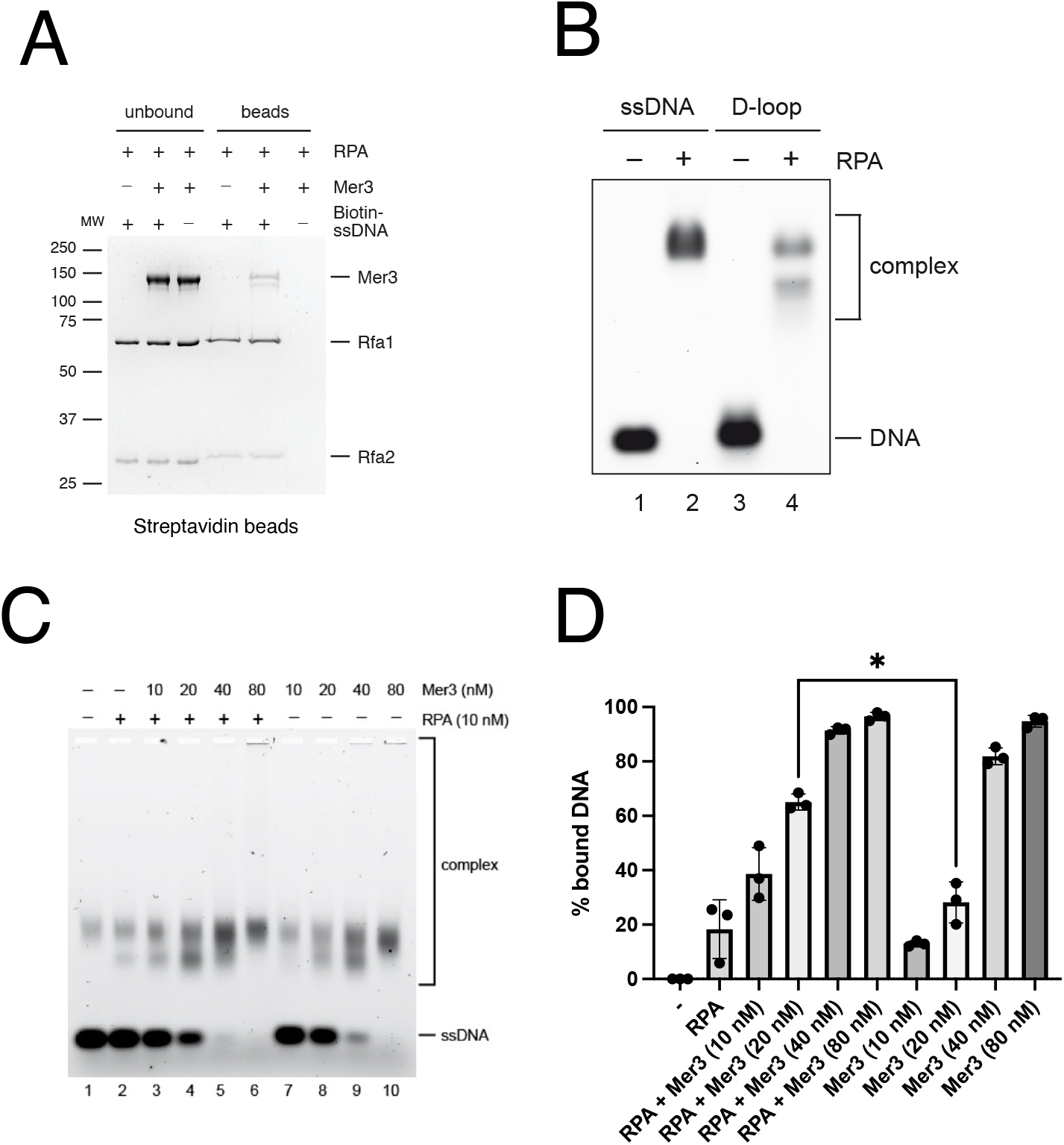
DNA binding properties of Mer3-RPA complex. A) Pulldown with 5^*′*^biotinylated 90-mer ssDNA (oWL1500) in the presence of RPA or Mer3 and RPA. B) EMSA showing the complexes formed with a constant concentration of RPA and an increasing concentration of Mer3 (left side) or an increasing concentration of Mer3 alone (right).

**Supplementary Figure 4.**
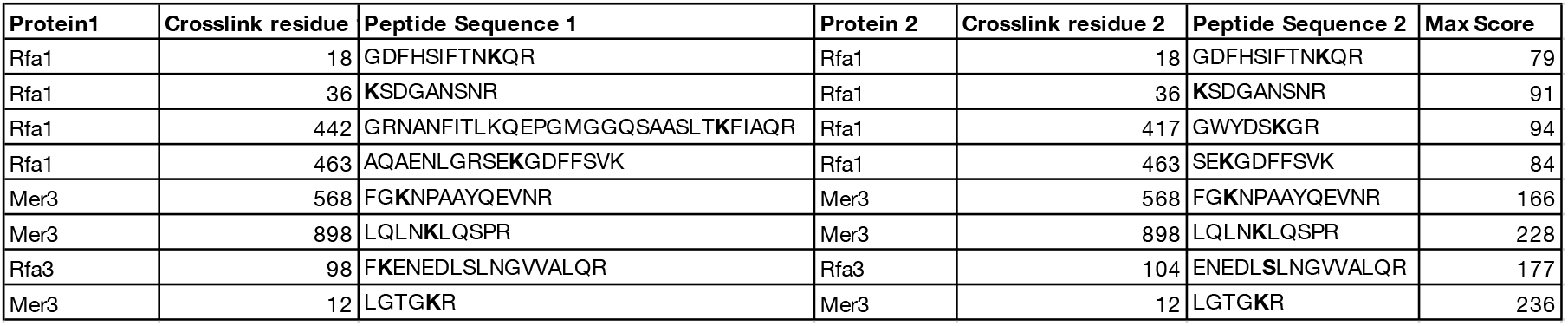
Summary of self-crosslinks in the Mer3-RPA complex.

**Supplementary Figure 5.**
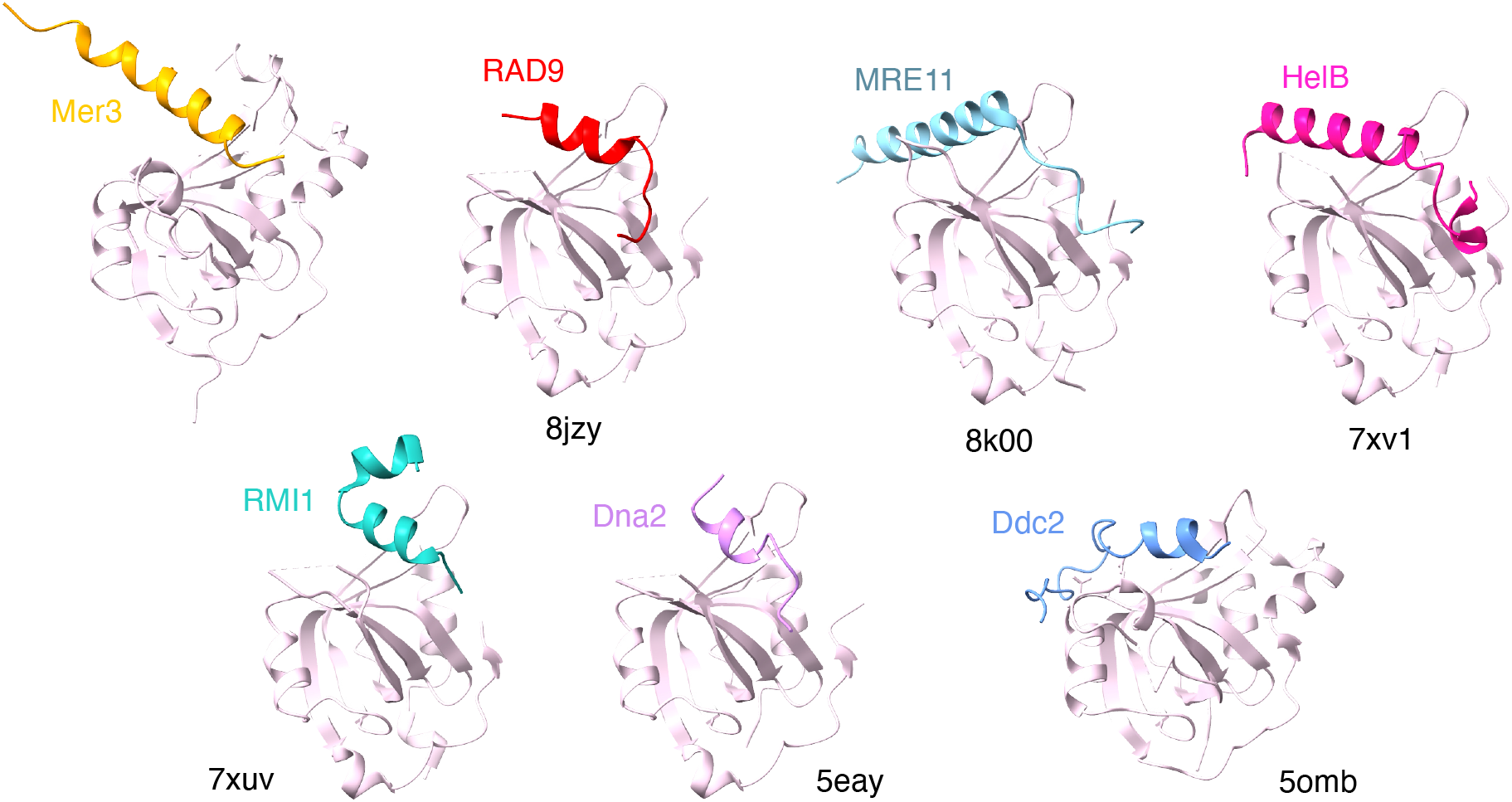
Structural comparison of N-OB binders. Structures of the N-OB binding helices are shown with the PDB IDs indicated. N-OB is shown in pale pink.

**Supplementary Figure 6.**
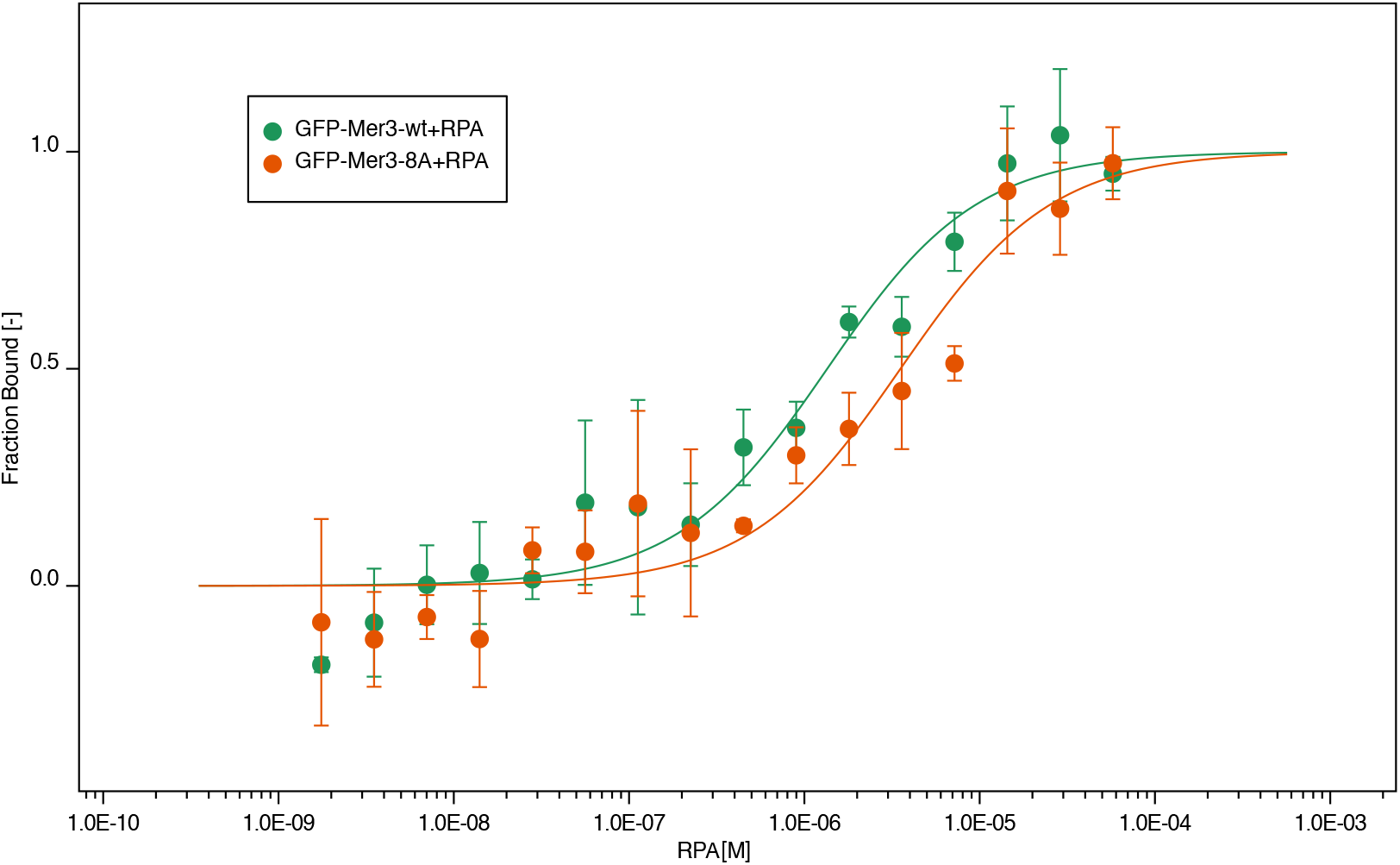
MST of Mer3 vs. Mer3-8A with RPA. Microscale thermophoresis (MST) of RPA binding to Mer3 wild type (green) or Mer3-8A mutant (orange). Increasing concentration of His-tagged RPA was titrated against N-terminally GFP-tagged Mer3 protein. Experiments were carried out in triplicate and approximate Kd of 1.31 μM +/-0.85 for Mer3-WT and 3.49 μM +/-1.03 for Mer3-8A was determined from the fitting curve in the NanoTemper Affinity Analysis v2.3 software (NanoTemper Technologies GmbH).

**Supplementary Figure 7.**
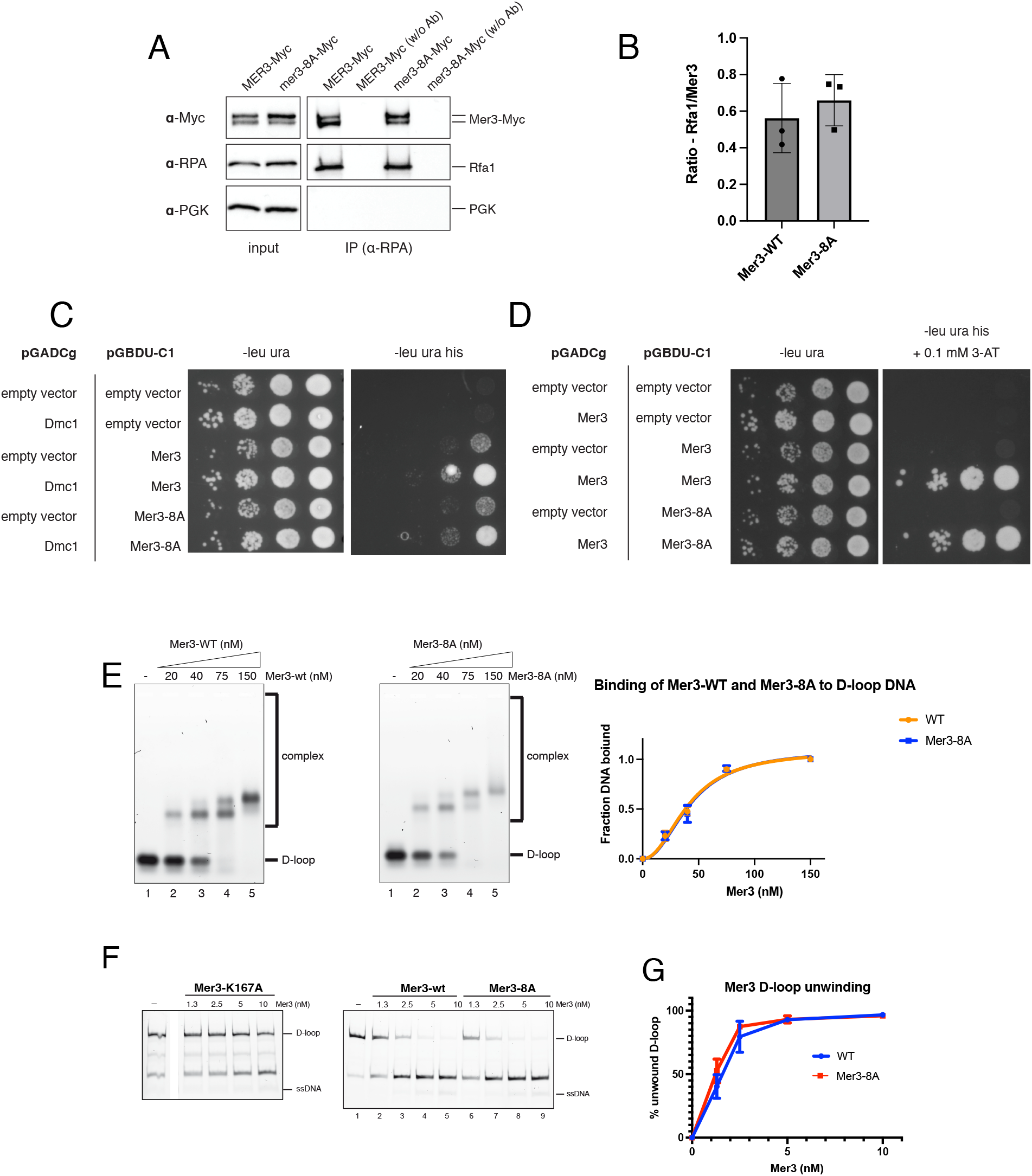
Mer3-8A retains all other binding and helicase properties as wild-type Mer3. A) co-IP of Mer3-9xMyc and Mer3-8A-9xMyc with RPA. B) Quantification of triplicate experiments as in A) C) Y2H showing that both Mer3 and Mer3-8A interact with Dmc1 recombinase, as previously shown [32]. D) Y2H showing that both Mer3 and Mer3-8A interact with itself, as previously shown [32]. E) EMSAs showing equivalent ssDNA binding properties for Mer3-WT (left) and Mer3-8A (right). F) Strand separation assays for Mer3 WT (middle), Mer3-8A (right) and the catalytically deficient Mer3-K167A (left). G) Quantification of three independent experiments shown in D).

**Supplementary Figure 8.**
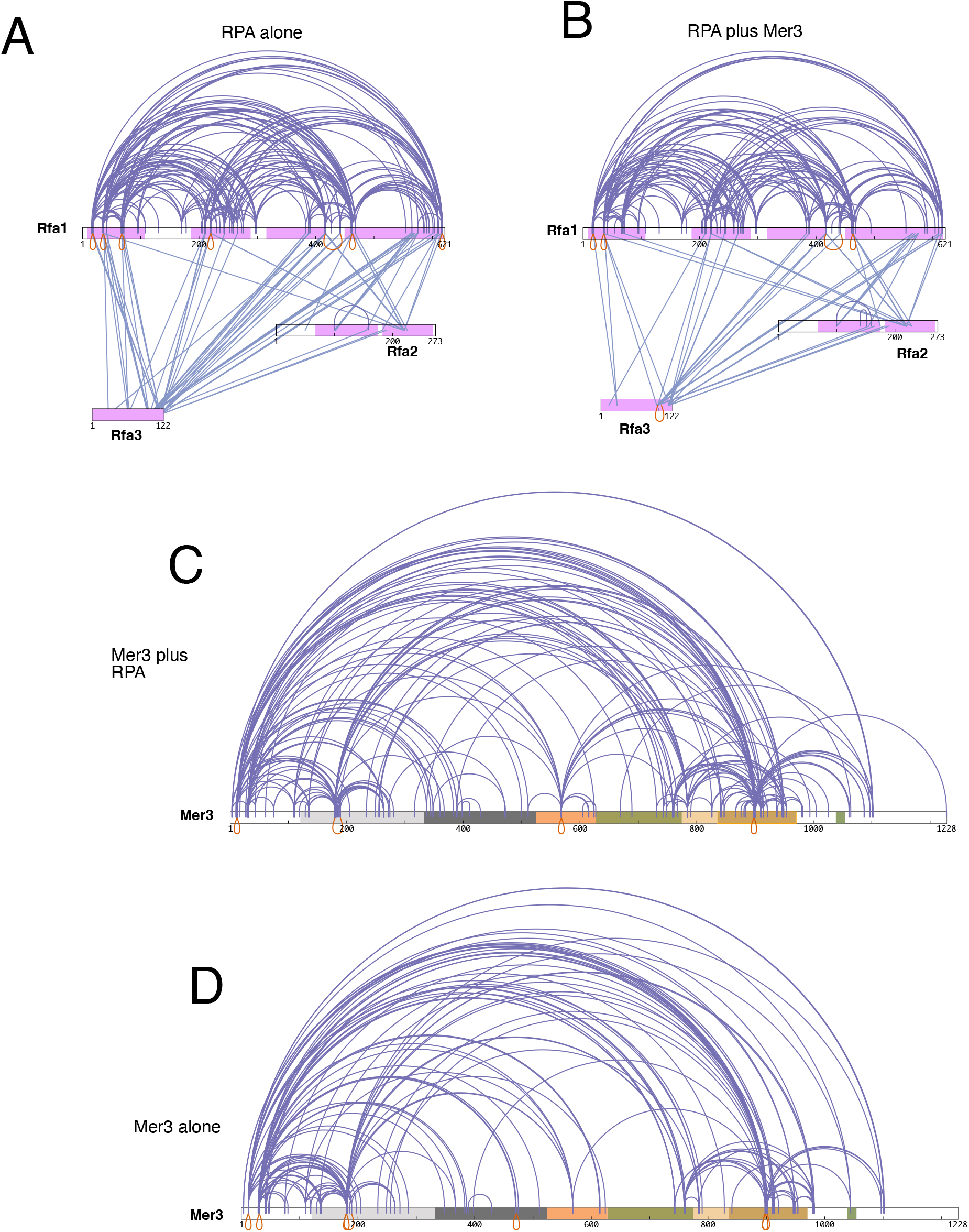
Comparison of XL-MS maps. A) Crosslinking map for RPA alone B) Crosslinking map for RPA in the presence of Mer3 C) Crosslinking map for Mer3 in the presence of RPA D) Crossliking map for Mer3 alone (taken from [32])

**Supplementary Figure 9.**
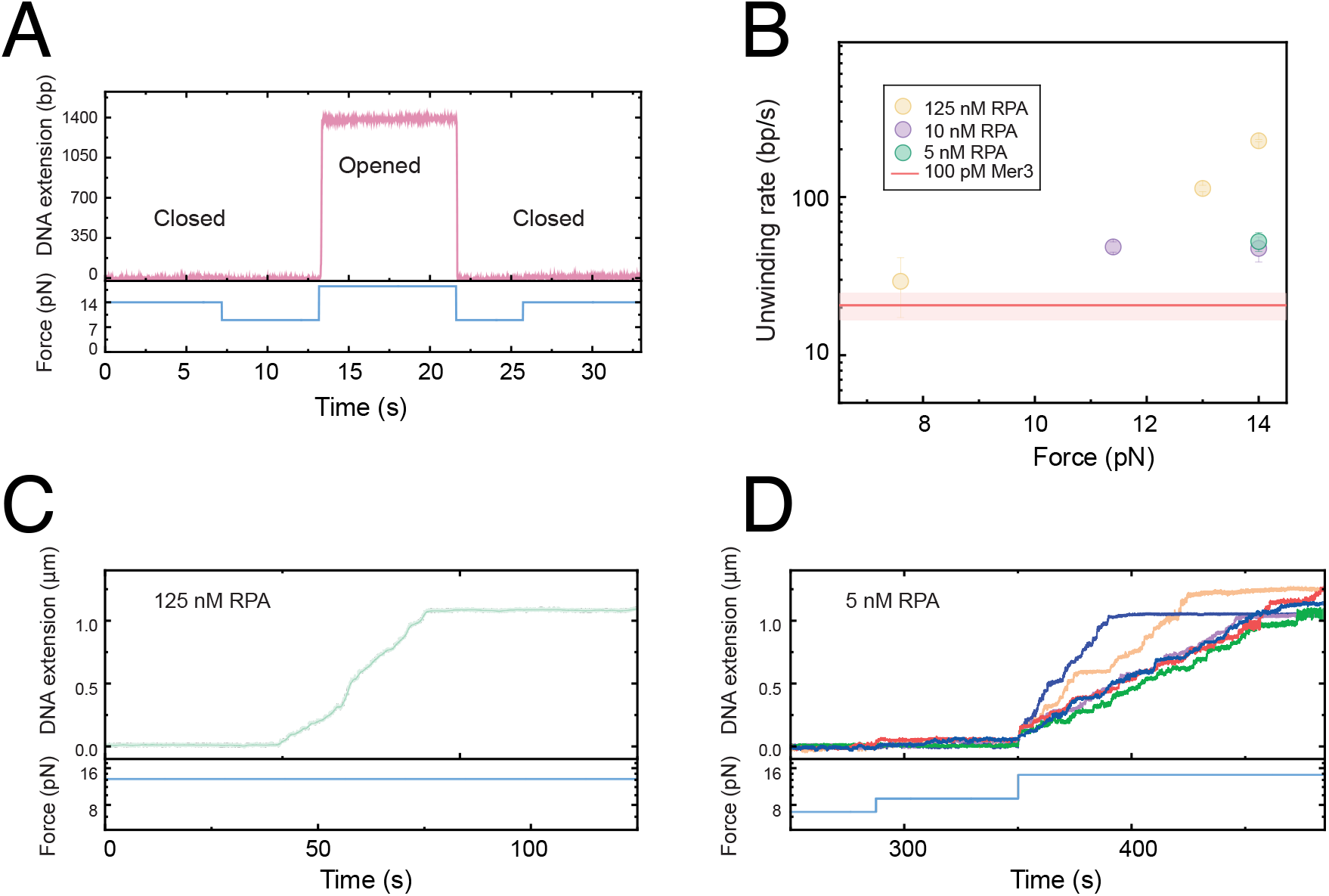
Magnetic Tweezer Single Traces. A) Validation of the hairpin DNA substrate: Application of force exceeding 15pN results in mechanical opening of the hairpin, observed as an increase in DNA extension due to the conversion of double-stranded DNA to single-stranded DNA. B) Mean pause-free unwinding rates by RPA as a function of applied force and RPA concentration. Significant force-dependent unwinding is observed primarily at RPA concentrations >10nM. The red shaded area represents the mean ± SD of the force-independent unwinding rate of Mer3 for comparison. C) Representative trace of hairpin unwinding by 125nM RPA at 14pN. D) Representative trace of hairpin unwinding by 5nM RPA only at 14pN. At lower forces, no unwinding activity was observed.

**Supplementary Figure 10.**
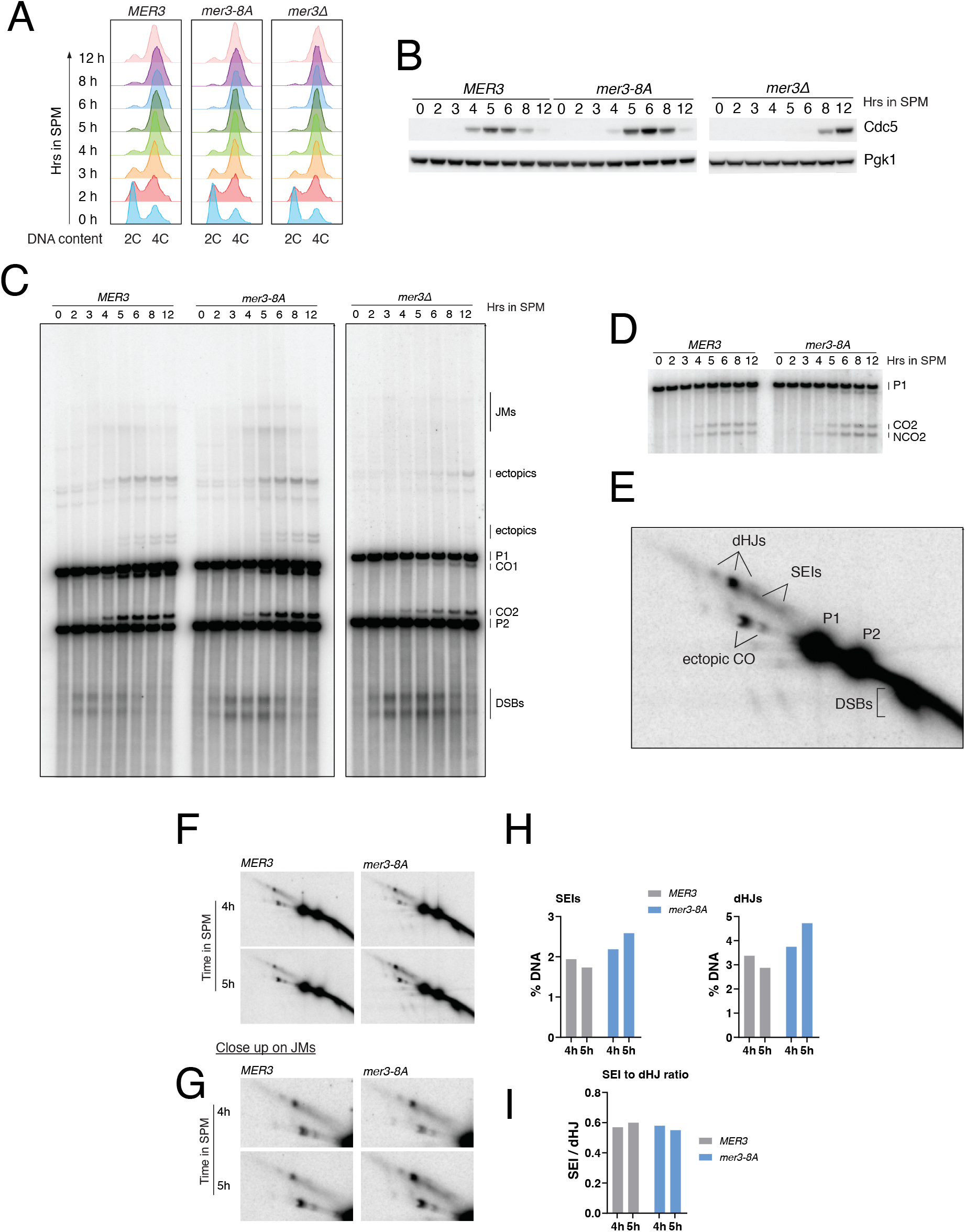
A) DNA content of the cells from a biological replicate of the experiment in Figure 5A, with the addition of data from a mer3 Δ strain. B) Cdc5 protein expression analysis from cells in (A). C) Physical analysis of recombination at HIS4-LEU2 in cells from (A). JMs – joint molecules, P1 and P2 – parental DNA, CO1 and CO2 – reciprocal recombinants of P1 and P2, DSBs – double-strand breaks, asterisk represents ectopic crossing over. Analysis of noncrossover and crossover formation at HIS4-LEU2 in cells from (A). D) Representative image of recombination intermediates at HIS4-LEU2. Psoralen-crosslinked DNA is digested with XhoI as in C) and analyzed by two-dimensional native/native gel electrophoresis followed by Southern blotting. E) Representative image of two-dimensional analysis of recombination intermediates showing what types of intermediates can be identified (dHJs - double Holliday junctions, SEIs - single end invasions, P1 and P2 – parental DNA, DSBs - double stranded DNA breaks F) Two-dimensional analysis of recombination intermediates at HIS4-LEU2 from cells in A). Timepoints with highest levels of JMs were selected based on one-dimensional analysis in C). G) Close-up images from F). G) Quantification of single-end intermediates (SEIs) and double Holliday junctions (dHJs) from E) and F), plotted as percentage of total DNA. H) SEI to dHJ ratio calculated from data in G).

**Supplementary Figure 11.**
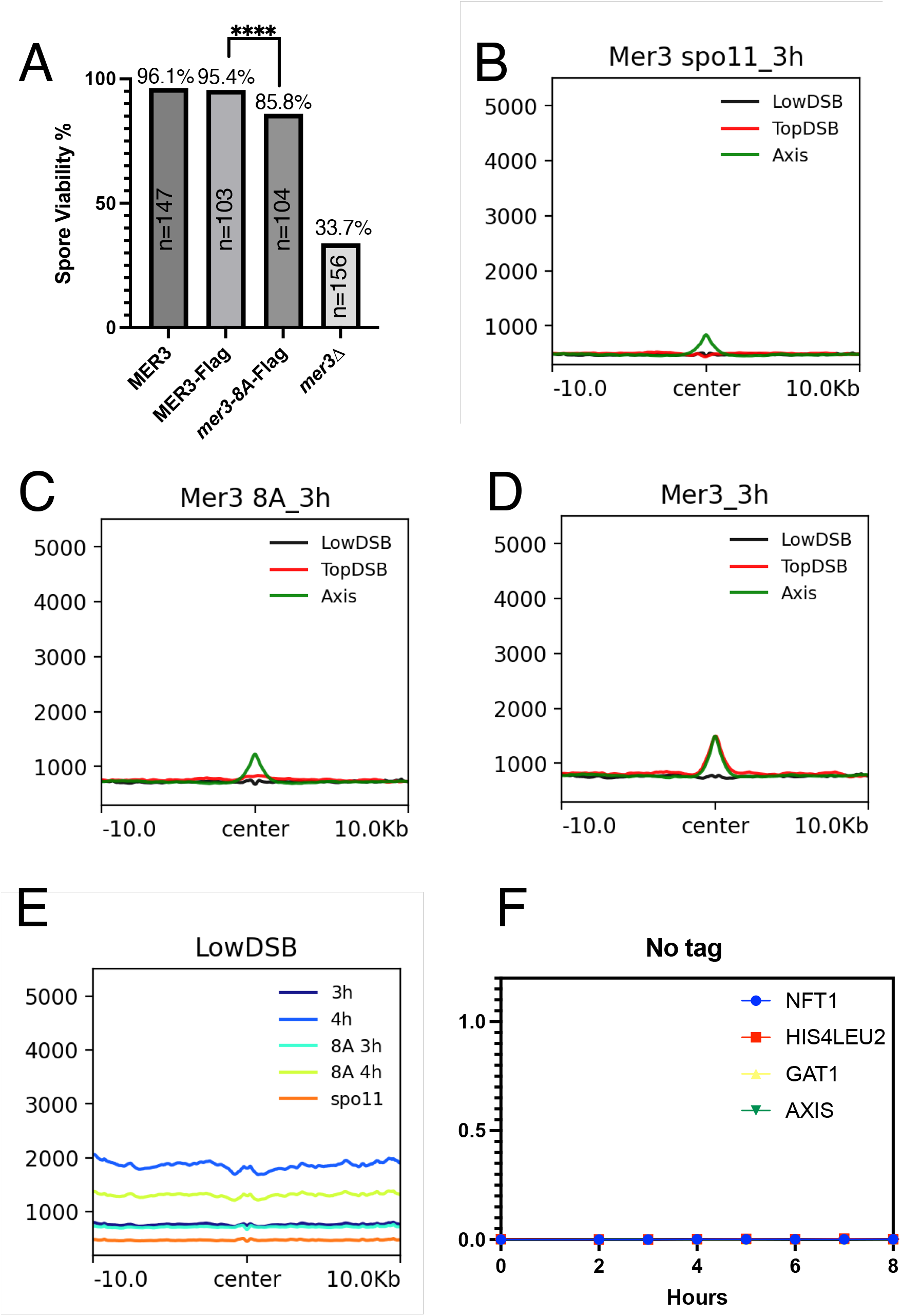
A) Spore viability for the Flag-tagged Mer3 strains. Number of tetrads dissected is indicated. Significance between MER3-Flag and mer3-8A-Flag determined by Fishers exact test (p= <0.001) B) localisation of Mer3-Flag in the Mer3-Flag/spo11Δstrain at the weakest 500 DSB sites (black), the strongest 500 DSB sites (red) and the axis (green) after 3 hours in sporulation media. C) Localisation of Mer3-8A-Flag in the mer3-8A strain at the weakest 500 DSB sites (black), the strongest 500 DSB sites (red) and the axis (green) after 3 hours in sporulation media. D) C) Localisation of Mer3-Flag in the MER3-Flag strain at the weakest 500 DSB sites (black), the strongest 500 DSB sites (red) and the axis (green) after 3 hours in sporulation media. E) Summary of localisation of Mer3-Flag or Mer3-8A-Flag in the backgrounds indicated for the weakest 500 DSB sites. F) Negative control for ChIP-qPCR experiment for Figure 5F and G.

